# Recording site placement on planar silicon-based probes affects neural signal quality: edge sites enhance acute recording performance

**DOI:** 10.1101/2020.06.01.127308

**Authors:** Richárd Fiáth, Domokos Meszéna, Mihály Boda, Péter Barthó, Patrick Ruther, István Ulbert

## Abstract

**Objective:** Multisite, silicon-based probes are widely used tools to record the electrical activity of neuronal populations. Several physical features of these devices (e.g. shank thickness, tip geometry) are designed to improve their recording performance. Here, our goal was to investigate whether the position of recording sites on the silicon shank might affect the quality of the recorded neural signal in acute experiments.

**Approach:** Neural recordings obtained with five different types of high-density, single-shank, planar silicon probes from anesthetized rats were analyzed. Wideband data were filtered (500 - 5000 Hz) to extract spiking activity, then various quantitative properties (e.g. amplitude distribution of the filtered potential, single unit yield) of the recorded cortical and thalamic activity were compared between sites located at different positions of the silicon shank, focusing particularly on edge and center sites.

**Main results:** Edge sites outperformed center sites: mean values of the examined properties of the spiking activity were in most cases higher for edge sites (~94%, 33/35) and a large fraction of these differences were also statistically significant (~45%, 15/33) with effect sizes ranging from small to large. Although the single unit yield was similar between site positions, the difference in signal quality was remarkable in the range corresponding to high-amplitude spikes. Furthermore, the advantage of edge sites slightly decreased for probes having a narrower shank.

**Significance:** The better signal quality on edge sites might be the result of the reduced shielding effect of the silicon shank providing a larger field of view for edge sites to detect spikes, or the less tissue damage caused near the edges of the shank. Our results might aid the design of novel neural implants in enhancing their recording performance by identifying more efficient recording site placements.

## 1. Introduction

To investigate the biological mechanisms behind complex brain functions (e.g. memory, sensation or consciousness) we have to monitor and manipulate the dynamics of neural populations of a statistically representative size, comprising hundreds or even thousands of neurons [1, 2]. Current research methods able to record the activity of so many individual cells simultaneously are extracellular electrophysiological recordings [3–8] and imaging techniques (e.g. two-photon calcium imaging, [7, 9–11]). Although most brain imaging approaches provide high spatial resolution, due to the slow kinetics of calcium indicators and limitations in the imaging speed, their temporal resolution is often low compared to electrophysiological techniques [12, 13]. This might hinder the precise registration of the action potential firing times of single cells, especially if the activity of many neurons is measured simultaneously. In contrast, recently developed high-density silicon-based multielectrode arrays [3–6, 14–19] can monitor the activity of hundreds of neurons at the same time with sub-millisecond precision and, because of the large number of closely packed recording sites, also with a high spatial resolution.

Planar silicon-based neural probes, which currently are among the most frequently used microelectrode types applied to measure extracellular brain activity, are fabricated using microelectromechanical system (MEMS) and complementary metal-oxide-semiconductor (CMOS) microtechnology. These methods allow to precisely realize the physical and geometrical parameters of these devices as well as to add integrated circuits on the probe shank or base for on-chip signal conditioning [20]. Considering the findings of previous research using neural implants for acute and chronic electrophysiological recordings, several features of silicon microprobes are now designed to minimize the mechanical trauma caused during probe insertion as well as to enhance their recording performance, both for the short and long term. For instance, in earlier studies it has been shown that the shank size of the neural implant has a significant impact on the extent of tissue damage caused by the insertion or on chronic tissue response [21–23]. Furthermore, the shape of the probe tip or the implant tethering may also affect the quality of the recorded neural signals in the long term [24–26]. Besides physical features of probes, the conditions of probe insertion (e.g. insertion speed, alignment of the probe) might also have a notably effect on the recording performance [24, 27–29].

An essential part of neural probes are the small recording sites which detect the electrical activity of neurons and are placed commonly on one of the sides of the silicon shank. The influence of size, impedance and material of recording sites on neural recordings is relatively well studied [30–33]. However, prior research investigating the optimal placement of recording sites on silicon shanks to achieve high quality recordings is scarce and contradictory [34–36]. In chronic experiments, Lee and colleagues found that recording sites placed on the edge of planar silicon probes perform slightly better than center sites [35]. The difference in signal quality was significant for wide (249 μm) devices but smaller and not significant for narrower (132 μm) probes. In contrast, the study of Scott and colleagues found no significant effect of site position on the recording quality in acute experiments [34]. However, in their study they only used data from one type of silicon probe (having an 85-μm-wide shank) and did not analyze single unit activity. Evaluating data obtained with a probe having a special edge electrode design showed a higher single unit yield and larger signal amplitudes on edge electrodes compared to sites located on the front side of the silicon shank [36]. Although edge sites outperformed the recording performance of other site positions, only a small number of recordings were analyzed in the study and signal amplitudes were only qualitatively compared.

As silicon probes and electrophysiological recording devices become affordable by more and more labs, which process is further accelerated by the open source movement [37–40], it is an important mission to thoroughly study various features of neural probes as well as brain-implant interactions to aid the design of future devices with the goal of enhancing their recording capabilities. An optimal placement of recording sites on planar silicon probes might increase the signal quality and the single unit yield significantly, and thereby decrease the costs and time requirements of experiments. Recording sites on commercially available passive silicon probes are usually placed in the center of the shank, although several variants of probes with edge sites also exist [41]. Contrary to that, recently developed silicon probes with high electrode density (e.g. the Neuropixels probe) contain sites both near the edge and in the center of their shank [3, 4, 14, 18, 42]. This site configuration provides an excellent opportunity to compare the recording performance of various site positions. In this comprehensive quantitative study, our aim was to examine whether the location of recording sites on high-density, single-shank, planar silicon-based probes might affect the quality of the recorded neural activity on acute timescales. In order to investigate this question, we examined electrophysiological recordings obtained with multiple probe types having different shank widths from the neocortex of anesthetized rats. Recording sites were labeled as edge or center sites according to their position on the silicon shank, then we analyzed the amplitude distribution of the recorded signal as well as various properties of single unit activity (e.g. single unit yield, peak-to-peak amplitude of single unit spike waveform) separately on these two site groups.

## 2. Methods

### 2.1 Silicon probe types

Neural recordings obtained with five different types of silicon-based, single-shank planar probes were analyzed in this study ([3, 14, 42]; figure 1). These probes have different shank widths (ranging from 50 μm to 125 μm), shank thicknesses (from 20 μm to 50 μm) and recording site features. All selected probe types have a high number of closely packed recording sites, ranging from 32 to 960 sites. To be able to make a reliable comparison in terms of site position, we only chose probes that contained recording sites both near the edge and the center of their silicon shank. Details of the features of these probes are listed below.

**Figure 1.**
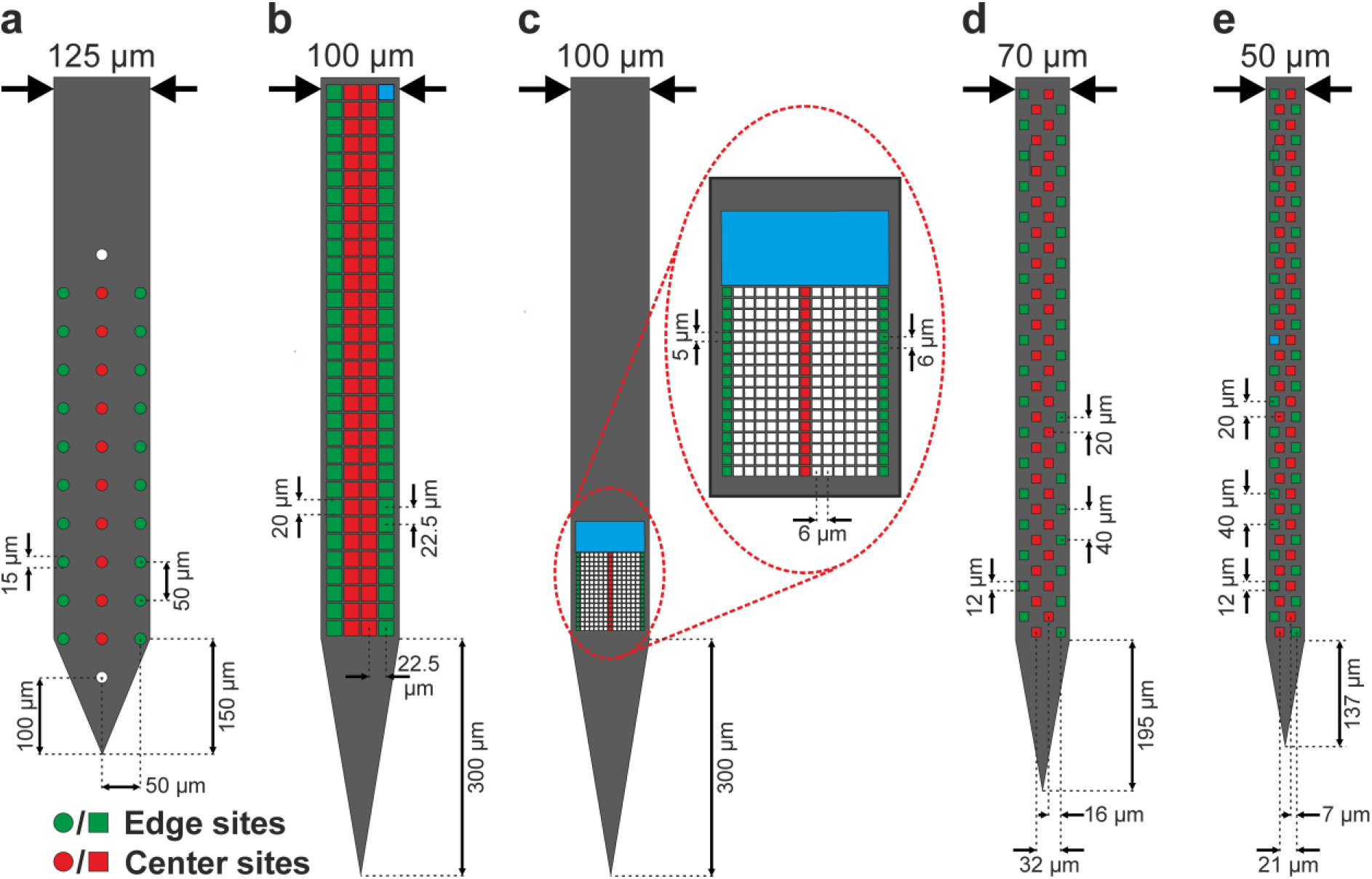
Schematic illustration of the five probe types examined in this study ordered by shank width. (a) 32-channel silicon polytrode (NeuroNexus). (b) 128-channel silicon probe (NeuroSeeker). (c) 255-channel silicon probe (NeuroSeeker). (d) Neuropixels probe with 70 μm shank width. (e) Neuropixels probe with 50 μm shank width. Recording sites classified as edge sites are colored green, while center sites are marked with red color. Sites colored white were either not located at the investigated positions or the data obtained by these were not used in the analysis. Blue sites on panel (b), (c) and (e) correspond to internal reference electrodes. Only a subset of recording sites is shown in the case of the Neuropixels probes.

The device with the lowest channel count was a commercially available silicon probe (A1×32-Poly3-10mm-50-177; NeuroNexus Technologies; www.neuronexus.com) with 32 iridium microelectrodes having a diameter of 15 μm (site area: 177 μm^2^), and a center-to-center distance of 50 μm (figure 1(a)). The silicon shank of the probe is 10 mm long and has a cross-section of 125 μm × 50 μm (Width (W) × Thickness (T)) at the level of the recording sites. The circular sites are arranged in three equidistantly spaced columns with one column located on each side of the shank (edge sites) and the third column in the center (center sites). The edge columns contain 10 microelectrodes each while the center column comprises 12 sites. The brain area covered by the recording sites is approximately 125 μm × 550 μm (W × Length (L)). The lowermost site (center column) is located 100 μm far from the probe tip. The closest point of edge sites from the edge of the silicon shank is 5 μm. This probe type is frequently used to record spiking activity from rats and mice [27, 43–45], was applied as mesh model in a computational modeling study [46], and was also used in the evaluation of the effect of site impedance on neural data quality [30].

The second device was a 128-site silicon probe with closely-spaced low-impedance titanium nitride electrodes recently developed in the NeuroSeeker research project ([14]; www.neuroseeker.com; figure 1(b)). This type of multielectrode has an 8 mm long shank with a cross-section of 100 μm × 50 μm (W × T). The spacing between the edges of the square-shaped recording sites (20 μm × 20 μm; site area: 400 μm^2^) is 2.5 μm. The sites are arranged in four equidistantly spaced columns (one column on each side and two columns in the center of the silicon shank) with all columns containing 32 microelectrodes. One microelectrode located at the top row on the right side has a larger are and serves as an internal reference electrode (only partially shown in figure 1(b)). The bottom row of recording sites is located 300 μm far from the chisel-shaped probe tip. The array of microelectrodes covers an area of 87.5 μm × 717.5 μm (W × L). The side of edge sites is approximately 6.25 μm far from the edge of the silicon shank. The 128-channel probe provides high-quality neural signals in acute experiments both from rodents and cats [14, 27, 43, 47], and was also chronically implanted in monkeys [48].

The third probe type was also developed in the NeuroSeeker project using the same fabrication technology (figure 1(c)). It has 255 miniature quadratic titanium nitride recording sites (5 μm × 5 μm; site area: 25 μm^2^). From this probe type, two different designs were fabricated, one with a linearly placed microelectrode array [49], and another with a closely-packed array of 17 × 15 recording sites [42]. The latter version was used in this study. The silicon shank of this probe has the same parameters as described above for the 128-channel probe. The spacing between the edge of small recording sites is 1 μm (center-to-center electrode distance is 6 μm). The probe has a large internal reference site located above the small sites. The bottom row of microelectrodes is located 300 μm from the chisel-shaped probe tip. The array of microelectrodes covers an area of 89 μm × 101 μm (W × L).

The fourth and fifth devices are two variants of the recently developed Neuropixels CMOS-based silicon probe ([3, 50]; www.neuropixels.org; figure 1(d) and (e)). The commercially available version has a 10 mm long shank and a cross-section of 70 μm × 20 μm (W × T; figure 1(d)). It contains 960 square-shaped titanium nitride microelectrodes (12 μm × 12 μm; site area: 144 μm^2^) from which 384 can be selected for recording. The recording sites are arranged in a checkerboard pattern with 4 columns and 480 rows. The center-to-center distance of microelectrodes in a single row is 32 μm. Alternate columns are offset by 16 μm and the vertical spacing of microelectrodes is 40 μm. The gap between the edge of the probe shank and the edge of the first recording sites is 5 μm. The center of the bottom row of sites is 195 μm away from the tip of the shank. The array of 384 adjacent recording sites covers a brain area of approximately 60 μm × 3800 μm (W × L). In addition, we analyzed data from a publicly available dataset obtained with another Neuropixels probe variant (PhaseA Option 1 probe; [50]). This probe has a shorter shank (5 mm), a smaller shank width (50 μm) and only 384 recording sites arranged in 4 columns and 192 rows (figure 1(e)). The features of recording sites are the same as described above. The center-to-center distance of microelectrodes in a single row is 21 μm and alternate columns are offset by 7 μm. The vertical spacing of microelectrodes is 40 μm and the side of edge sites is located 5 μm from the edge of the silicon shank. The center of the bottom row of sites is 137 μm away from the tip of the shank. The brain area covered by the recording sites is approximately 40 μm × 3800 μm (W × L). Neuropixels probes are being increasingly used in electrophysiology labs and became essential tools of cutting-edge neuroscience research [6, 51, 52].

The probes investigated in this study were principally designed for acute *in vivo* recordings, although Neuropixels probes can be used for chronic experiments [53, 54] and the 32-channel NeuroNexus probe is also available with a chronic design. Except for the NeuroNexus probe, all probes were fabricated using a 0.13-μm CMOS fabrication process. All probe types were passive devices, except for the Neuropixels probes which contain on-chip electronics for signal conditioning (e.g. for filtering, amplification or multiplexing) and digitization on the probe [3]. Electrical impedance of titanium nitride sites of CMOS-fabricated probes was low (< 1 MΩ at 1 kHz) but varied between different probes types due to the difference in the area of recording sites. However, the absolute impedance magnitude values of a particular probe type showed low variability (about a few kΩ; [3, 14, 49]). The site impedance of the NeuroNexus probe measured at 1 kHz was 385.63 kΩ ± 30.7 kΩ (mean ± standard deviation; average of 64 recording sites of two probes). No statistically significant difference was found between the impedance of edge and center sites of the examined probe types.

### 2.2 Analyzed datasets

To obtain data for analysis, we performed acute experiments in anesthetized rats and recorded spontaneously occurring cortical activity with each probe type (see sections 2.3 and 2.4 below for details), except for the Neuropixels probe with 50 μm shank width. In the latter case, recordings from a publicly available online database were used for analysis (n = 7 out of 43 files: c5, c8, c12, c24, c26, c32, c45; [50]). All public Neuropixels data files were visually inspected to select recordings with the best quality. To avoid data redundancy, only one recording file was used from a dataset containing multiple recordings which were measured from the same penetration. A subset of the 128-channel probe data analyzed here originated from the dataset obtained in another study [27]. Details of the recordings per probe type can be found in table 1.

**Table 1.** Details of the experiments and recordings of the five silicon probe types.

| Probe type | No. of recording sites | No. of channels in separated recordings | No. of analyzed recordings | No. of rats | No. of penetrations | Insertion speed (mm/sec) | Type of anesthesia | Average recording length (min) | Reference |
| --- | --- | --- | --- | --- | --- | --- | --- | --- | --- |
| 32-channel NeuroNexus probe | 32 | 10 | 10 | 6 | 10 | 0.002 | Ketamine/xylazine | 31 | www.neuronexus.com |
| 128-channel NeuroSeeker probe | 128 | 32 | 10 | 10 | 10 | 0.002 | Ketamine/xylazine | 30 | [14] |
| Neuropixels probe (70 $\mu\text{m}$ wide) | 384 | 58 | 6 | 4 | 6 | ~1 | Urethane | 31 | www.neuropixels.org |
| Neuropixels probe (50 $\mu\text{m}$ wide) | 384 | 58 | 7 | 6 | 7 | 0.005 | Urethane | 20 | [50] |
| 255-channel NeuroSeeker probe | 255 | 17 | 21 | 2 | 2 | 0.002 | Ketamine/xylazine | 32 | [42, 49] |

Publicly available cortical recordings (n = 7) acquired by the 255-channel NeuroSeeker probe were also analyzed in this work (recordings used: Co1-Co3, Co5 and CoP1-CoP3; www.kampff-lab.org/ultra-dense-survey; supplementary table 1; [42]). However, because of differences in the experimental conditions and data quality, these public recordings and our 255-channel measurements were assessed separately.

To examine the effect of sample size on the results, a larger cortical dataset acquired during previous projects (e.g. [14, 27]) with the 128-channel probe was also included for analysis. A total of 186 recordings (n = 41 rats and ~100 penetrations) with durations ranging from 5 to 45 minutes were examined. The recordings contained activity from each layer of the rat cortex.

To examine the differences in signal quality between edge and center sites in another brain structure, a small dataset obtained from the thalamus was also evaluated (n = 9 recordings; supplementary table 2).

### 2.3 Animal surgery

All experiments were performed according to the EC Council Directive of September 22, 2010 (2010/63/EU), and all procedures were reviewed and approved by the Animal Care Committee of the Research Centre for Natural Sciences and by the National Food Chain Safety Office of Hungary (license number: PEI/001/2290-11/2015). In the case of the 32-channel, 128-channel and 255-channel probe types, acute *in vivo* experiments were carried out similarly as described in our earlier studies [14, 27, 49]. In short, Wistar rats were anesthetized with a mixture of ketamine (75 mg/kg of body weight) and xylazine (10 mg/kg of body weight) injected intramuscularly. If necessary, supplementary ketamine/xylazine injections were given to maintain the depth of anesthesia during surgery and recordings. The animals were placed in a stereotaxic frame (David Kopf Instruments, Tujunga, CA, USA) after they reached the level of surgical anesthesia. The body temperature of rats was maintained with a homeothermic heating pad connected to a temperature controller (Supertech, Pécs, Hungary). After the removal of the skin and the connective tissue from the top of the skull, a craniotomy with a size of about 3 × 3 mm^2^ was drilled over the neocortical area of interest (trunk region of the somatosensory cortex (S1Tr); approximate coordinates of the target site were: anterior-posterior (AP): –2.7 mm; medial-lateral (ML): 2.5 mm; with respect to the bregma [55]). Then, to avoid excessive brain dimpling during the insertion of the single-shank silicon probes, a small slit was carefully made in the dura mater above the insertion site using a 30-gauge needle. For a post-mortem histological verification of the recording location of the probe [56], the silicon shank was coated with red-fluorescent dye 1,1-dioctadecyl-3,3,3,3-tetramethylindocarbocyanine perchlorate (DiI, D-282, ~1% in absolute ethanol, Thermo Fischer Scientific, Waltham, MA, USA) before insertion. After that, the silicon probe mounted on a motorized stereotaxic micromanipulator (Robot Stereotaxic, Neurostar, Tübingen, Germany) was driven into the brain tissue with a slow insertion speed of 2 μm/sec to decrease tissue damage [27]. During probe insertion, care was taken to avoid damaging blood vessels located on the brain surface. With the 32-channel and 128-channel probes, neural activity was recorded mostly from cortical layers where the neuronal spiking activity is the strongest during ketamine/xylazine-induced slow wave activity (layers IV-V; [57]). However, the depth of recording varied slightly between penetrations. With the 255-channel probe, because it records only from a confined cortical region (~100 μm × 100 μm), we acquired during a single penetration data from multiple depths (using at least 100 μm insertion steps to avoid recording form overlapping areas) from cortical layers where spiking activity could be detected (layers III-VI). In the case of thalamic recordings acquired with the 128-channel probe, the probe was moved below the neocortex to a dorsoventral depth of about 4.5 mm – 6.5 mm, to target somatosensory thalamic nuclei (the ventrobasal complex and the posterior nucleus).

Room temperature physiological saline solution was regularly dropped into the cavity of the craniotomy to prevent dehydration of the neocortex. A stainless steel needle inserted in the neck muscle of the animal served as the reference and ground electrode during recordings.

In the case of the experiments with the 70-μm-wide Neuropixels probe, the rats were anesthetized using urethane (1.5 g/kg). The experimental procedure was similar as described above. The probes were driven into the parietal association cortex (AP: −4.1 mm, ML: 3 mm; with respect to the bregma) by hand with a stereotaxic micromanipulator to a depth of ~3 mm, using an insertion speed of approximately 1 mm/sec. Details of experiments and recordings are summarized in table 1.

At the end of the experiments, probes were withdrawn and cleaned by immersing them into 1% Tergazyme solution (Sigma-Aldrich, St. Louis, MO, USA) for at least 30 minutes followed by rinsing with distilled water for about 2 minutes.

### 2.4 Electrophysiological recordings

For the three passive probe types, spontaneously occurring brain electrical activity was recorded using an Intan RHD-2000 electrophysiological recording system (Intan Technologies, Los Angeles, CA, USA). Two 128-channel amplifier boards were used in case of the 255-channel probe, one 64-channel and two 32-channel amplifier boards were used in the case of the 128-channel probe, and a single 32-channel board was used with the 32-channel probe. The recording system was connected to a laptop via USB 2.0. Wideband signals (0.1 – 7500 Hz) were recorded with a sampling frequency of 20 kHz/channel and a resolution of 16 bit. About 30 minutes of multichannel neuronal data were collected at a single recording location. Data were saved to a local network attached storage device for offline analysis.

In the case of the 32-channel probe, two probes were used for the experiments. One probe was implanted into the neocortex of five rats, while the other probe was used in one rat. One or two probe insertions were done in each animal (n = 10 penetrations in total). All recording sites of the probes were functional. Two identical probes from the same manufacturing batch were used during the experiments with the 128-channel probes. Each probe was implanted in five rats (n = 10 penetrations in total). The probes contained a maximum of two unfunctional recording sites. Two penetrations were carried out with the 255-channel probe (n = 2 rats). The used probe had multiple unfunctional recording sites (n = 14 sites; the site map is shown in supplementary figure 1(a)).

In the case of the Neuropixels probes, data recorded in the action potential band (AP, 300 – 10.000 Hz) was used. The sampling rate was 30 kHz/channel and digitization was performed at 10 bits, under a gain of 500, yielding a resolution of 2.34 μV per bit. Data was acquired using the SpikeGLX open-source software (github.com/billkarsh/SpikeGLX). The 70-μm-wide probe had one internal reference site, while the 50-μm-wide device had 12 internal reference electrodes.

### 2.5 Grouping of recording sites

Recording sites were either classified as edge or center sites based on their location on the silicon shank (figures 1 and 2). For the 32-channel probe, the two columns of sites located on the sides of the shank were labeled as edge sites, and two 10-channel recording files were created from the original 32-channel data (edge sites; figure 1(a)). By removing the top and bottom recording sites (to match the channel number of files containing edge channels), we constructed a 10-channel recording file from the middle column of 12 sites (center sites; figure 1(a)). For the 128-channel probe, four 32-channel recording files were generated from the original 128-channel data based on the location of the columns of sites (figure 1(b) and figure 2). Microelectrodes located on the sides were classified as edge sites while the sites in the middle two columns were categorized as center sites. For the 255-channel probe, the column of recording sites located on the left side of the probe were classified as edge sites, while the sites in the 8^th^ column were grouped as center sites, both containing 17 channels (figure 1(c)). The sites located on the right edge were not used in the analysis because of the high number of unfunctional channels (supplementary figure 1(a)). In the case of the public 255-channel dataset, both edge columns were included in the analysis, as the number of bad sites was low on the edges (supplementary figure 1(b)). For the Neuropixels probes, the assignment of site locations was similar as described for the 128-channel probe (figure 1(d) and (e)). After removing channels corresponding to reference electrodes and their neighbors, as well as recording sites located outside of the cortex, the constructed individual recording files had 58 channels each.

**Figure 2.**
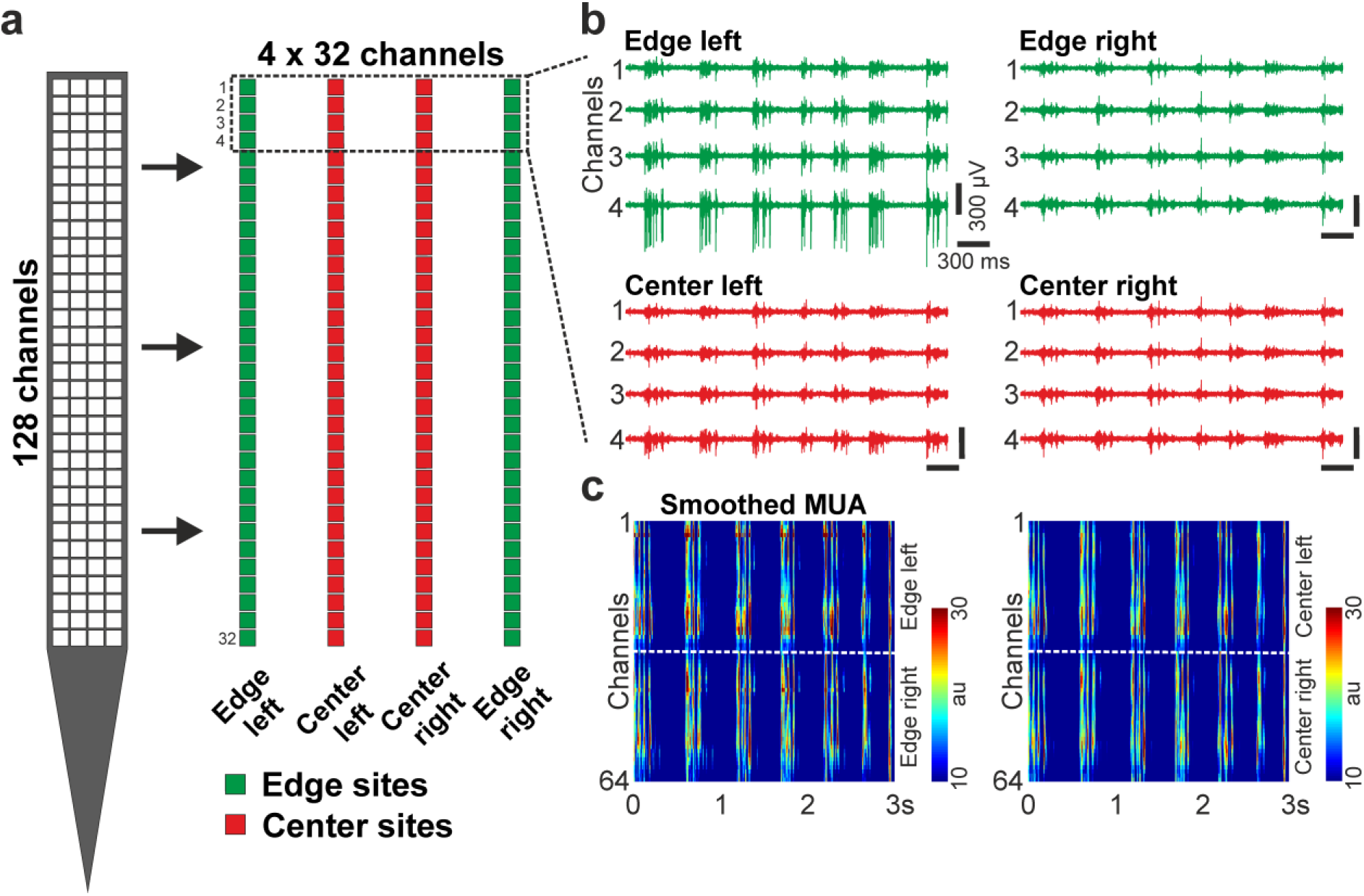
Grouping of edge (green) and center (red) sites. (a) Recordings obtained by the silicon probe (e.g. the 128-channel NeuroSeeker probe shown here) were separated into groups of channels (4 × 32 channels here) based on the position (edge or center) of recording sites. (b) Sample three-second-long multiunit activity (MUA; 500-5000 Hz) traces recorded on the channels indicated with a dashed rectangle on panel a. (c) Rectified and smoothed MUA (50 Hz lowpass filter) recorded on all edge (left) and center (right) channels. The dashed white lines separate channels located on the left and right side of the probe. au, arbitrary unit.

All analyses were performed separately on the new data files created based on the position of recording sites. Since there was no statistically significant difference in the signal quality between left and right sides of the probes (supplementary figures 7-10(a)-(e); recsupplementary tables 4-7), results of the analyses obtained on data files that belonged to the same site group (e.g. left edge and right edge sites) were pooled and are presented together. However, the supplementary material contains also the results obtained on all separate recordings (supplementary figures 7-10(a)-(e); supplementary tables 4-7).

In the case of the recordings obtained with the 128-channel probe, 32-channel files were generated also based on the longitudinal positions of the recording sites (supplementary figure 15(a)).

### 2.6 Amplitude distribution of the filtered potential

Amplitude histograms of the filtered potential (500 – 5000 Hz, Butterworth 3^rd^-order bandpass filter, zero-phase shift) were constructed to examine the difference in the signal amplitudes between edge and center sites (figure 3(a)). Channels corresponding to internal reference electrodes as well as bad sites were excluded from the analysis. Every 50^th^ sample on each channel was used to create the histogram (resulting in several millions of samples per site position). Then, in four non-overlapping amplitude ranges (figure 3(b)), we calculated the ratio of the number of samples in a particular amplitude range to the total number of samples examined in a site group, both for edge and center sites (figure 3(c)). The selected ranges were chosen, based on the features of the analyzed data and on our previous experience, to match the spike amplitudes of single units with different levels of quality (or signal-to-noise ratio): units with questionable quality (−50 μV to −100 μV), average quality units (−100 μV to −200 μV), high-quality units (−200 μV to −300 μV) and single units with superior quality (−300 μV to −1000 μV). Because spike-like events with amplitudes lower than −1000 μV are usually the results of artifacts, potential values below −1000 μV were not examined.

**Figure 3.**
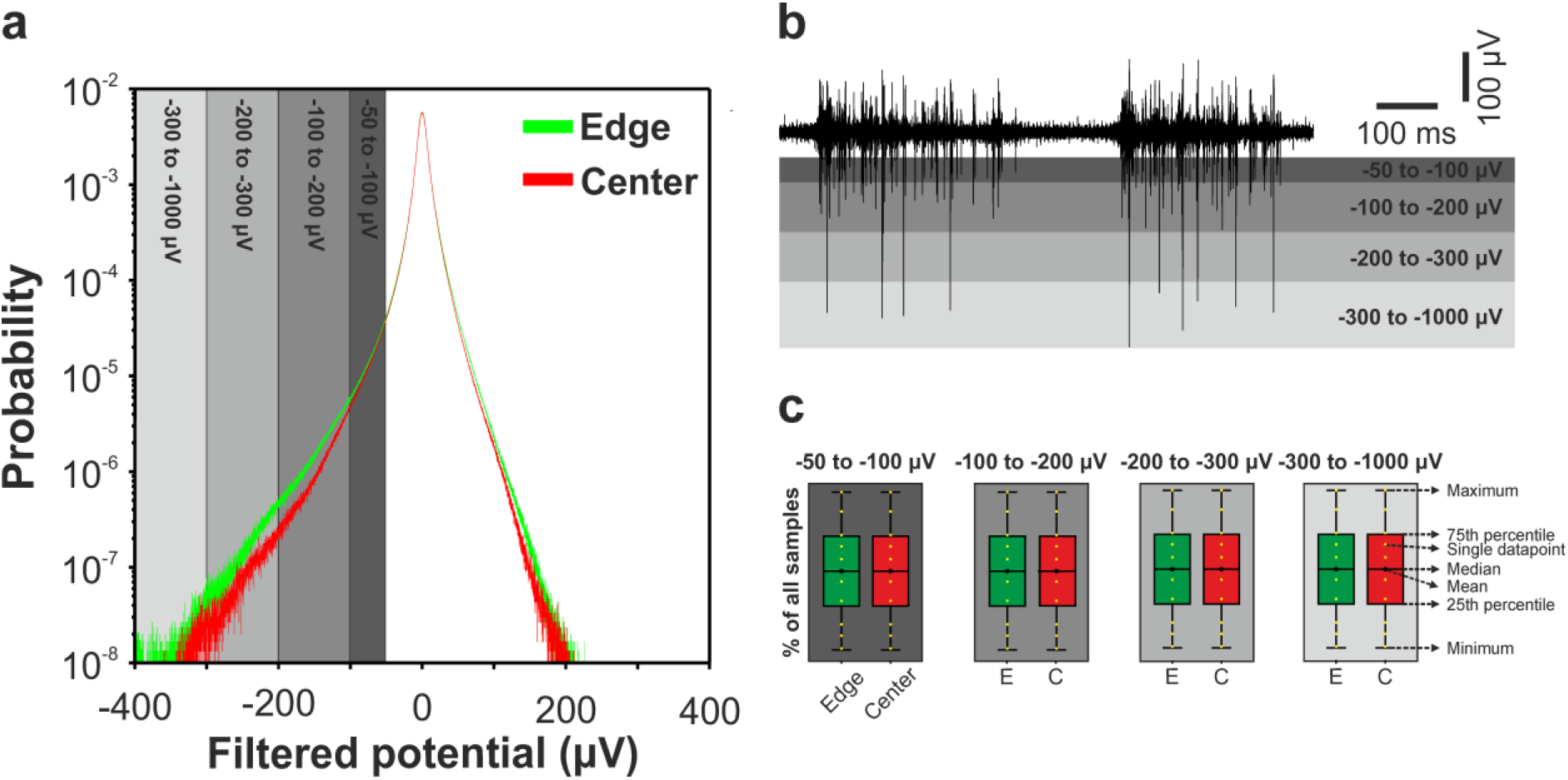
Amplitude distribution of filtered potentials. (a) Representative logarithmic histogram of sample values of a filtered (500-5000 Hz) recording obtained on edge (green) and center (red) sites of a 128-channel probe. The gray areas indicate the four amplitude ranges investigated in the study. (b) A single-channel, one-second-long filtered trace of the same recording superposed on the four amplitude ranges (gray rectangles). The lowermost range (light gray) is only partially shown. (c) To compare the amplitude distribution of edge (E) and center (C) sites, color-coded boxplots were constructed for each amplitude range of each silicon probe. Features of boxplots are indicated on the right side. The presented boxplots are only for demonstration and do not contain real data.

By evaluating the distribution of negative amplitudes only, we can describe the signal quality more precisely compared to the case when positive amplitudes are also included. The reason behind this is that, in referential recordings, extracellular spikes have usually a negative peak with a significantly larger amplitude compared to the amplitude of their positive peaks. Here we present results based on the analysis of negative amplitudes. However, the same analysis was performed also for positive amplitudes using the same amplitude ranges. Results of this analysis can be found in the supplementary material (panel (f) of supplementary figures 7-10; supplementary figure 11(a); supplementary tables 4-8).

### 2.7 Spike sorting and calculation of single unit properties

To assign the recorded spikes to individual neurons, automatic spike sorting was performed on recordings separated by site position using the MATLAB-based software Kilosort [58]. Manual revision of the single unit clusters detected by Kilosort was done in Phy, an open source neurophysiological data analysis package written in Python (github.com/kwikteam/phy). The manual revision was done blindly, that is, the user did not know whether the actual data file was recorded by edge or center sites. After the revision of spike sorting results, wideband spikes of each single unit cluster were averaged together to obtain the average spike waveforms. For further analysis, we selected well-isolated units using the following criteria [27]. We defined a single unit as well isolated if it had a clear refractory period (less than 2% of the spikes in the 2-ms-long refractory period), a firing rate higher than 0.05 Hz (or at least 100 spikes in the cluster) and a spike waveform with a peak-to-peak amplitude over 60 μV. The peak-to-peak amplitude was defined as the amplitude difference between the negative peak (or trough) and the largest positive peak of the average spike waveform, computed on the recording channel which contained the spikes of the particular single unit with the highest amplitude. These criteria allowed us to exclude low quality units as well as to decrease the effect of subjective decisions of the operator during the manual curation of neuron clusters. Only a low percent of the units was excluded from the analysis (from 1 to 12%; supplementary table (3)). The following single unit properties were calculated and used to compare the signal quality of edge and center sites: single unit yield, peak-to-peak amplitude of the average spike waveform of each well-separated single unit, and the isolation distance of each unit cluster (github.com/cortex-lab/sortingQuality; [59]).

### 2.8 Estimation of the noise level of the filtered signal in vivo

Although it would be possible to measure the noise level of recording sites using data obtained *in vitro* in saline solution, we do not had access to all probes types to perform these tests. Therefore, we developed a method to estimate the noise level based on *in vivo* recordings. Because most cortical neurons cease to fire for a couple of hundred milliseconds during down-states of the ketamine-xylazine or urethane-induced slow wave activity [57], the signals recorded during these short time windows of neuronal silence might be appropriate to approximate the noise level of recordings. We used a state detection algorithm previously developed by our group [57] to detect the onset of up- (high spiking activity) and down-states (low spiking activity; figure 4). First, the wideband signal was filtered (500 - 5000 Hz; Butterworth 3^rd^-order bandpass filter; zero-phase shift) and rectified to extract the multiunit activity (MUA). After that, all channels were summed up sample-wise (Summed MUA; figure 4) then smoothed using a 50 Hz lowpass filter to extract the envelope of the MUA (Smoothed MUA; figure 4). Next, using a threshold level (calculated by also taking into account the duration of slow wave states), we detected the state onsets. Finally, on each channel of the rectified MUA, the root mean square (RMS) value of a 50-ms-long segment in the middle of down-states with a duration of at least 200 ms was calculated, then the RMS values were averaged (figure 4). The method was validated by *in vitro* measurements of the RMS noise of recording sites of 128-channel and 255-channel probes in saline solution. For the 128-channel probe (n = 6 probes), the estimated noise level in the 500 – 5000 Hz frequency range was around 80% higher than the noise level measured *in vitro (in vivo* vs. *in vitro:* 3.87 ± 0.43 μV_RMS_ vs. 2.17 ± 0.66 μV_RMS_; mean ± standard deviation). For the 255-channel probe (n = 2 probes), this difference was about 65% (*in vivo* vs. *in vitro:* 6.87 ± 0.41 μV_RMS_ vs. 4.15 ± 0.58 μV_RMS_). A similar difference was also found in the case of the Neuropixels probes (10.22 ± 2.01 μV_RMS_ for the 50 μm wide probe and 8.45 ± 0.81 μV_RMS_ for the 70 μm wide probe vs. 5.1 ± 0.6 μV_RMS_ reported *in vitro* in [3]). Thus, although our method slightly overestimates the noise level, it is suitable to measure this property and compare it between edge and center sites.

**Figure 4.**
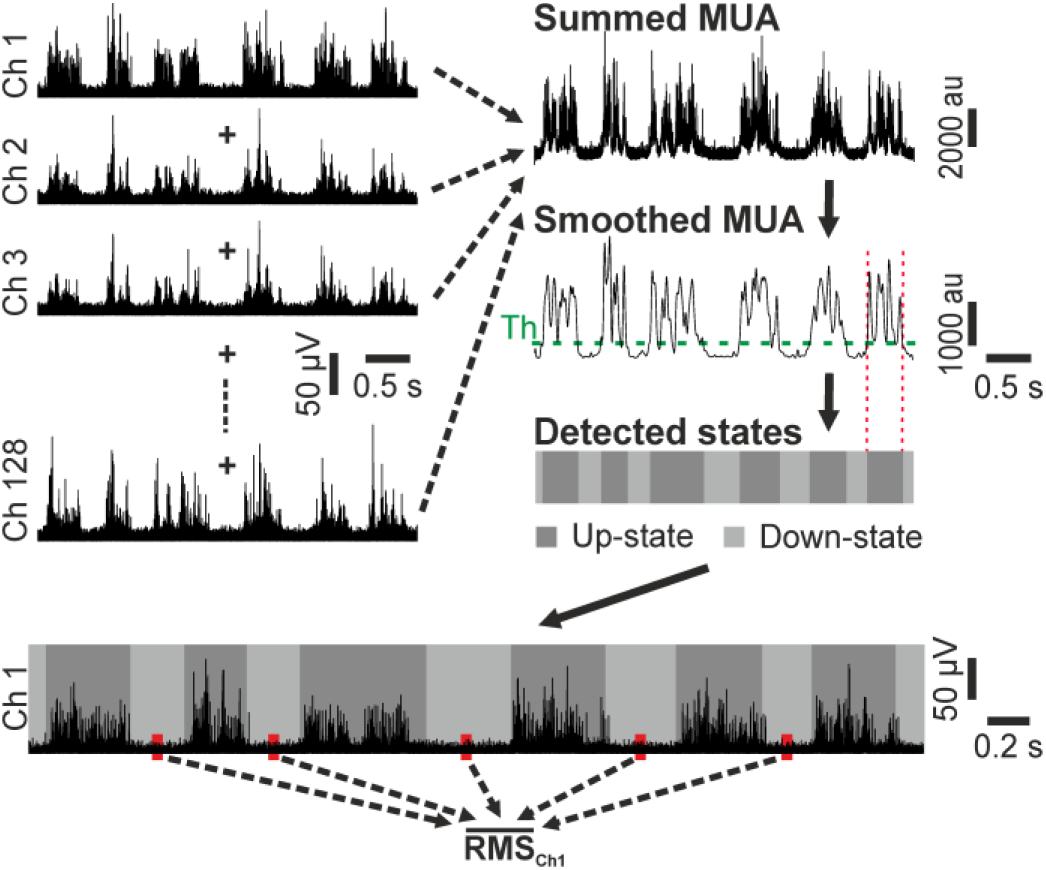
The method used to estimate the noise level of silicon probe recordings *in vivo*. The onsets of up- and down-states were detected based on the MUA, then the root mean square (RMS) values of short segments of recordings were calculated in the center of down-states (low spiking activity) and averaged for each channel. A detailed description of the noise estimation method can be found in the section 2.8.

### 2.9 Statistical analysis

Because most of the variables of interest had a low sample size or non-normal distribution (according to the Kolmogorov-Smirnov and Shapiro-Wilk tests) we used non-parametric tests for statistical analysis. For the amplitude distribution (matched samples), we used the Wilcoxon signed-rank test to compare the performance of edge and center sites (two groups), and the Friedman test was used to compare difference also between left and right sides (three or four groups). For other variables (noise level, single unit yield, unit amplitude and isolation distance), the Mann-Whitney U test was applied to compare the performance of edge and center sites (two groups), and Kruskal-Wallis test was used for the comparison of laterality (three or four groups). When a significant difference was found between site positions by the Friedman or the Kruskal-Wallis test, post-hoc analysis was performed for all pairwise comparisons using Dunn’s test with Bonferroni correction. p values less than 0.05 were considered significant. Effect sizes were calculated using the following formula: r = Z/√N, where Z is the z-score and N is the sample size. Features of boxplots (figures 3 and 5–9; supplementary figures 7-15) showing the distribution of data are presented as follows. The middle line indicates the median, while the boxes correspond to the 25th and 75th percentile. Whiskers mark the minimum and maximum values. The average is depicted with a black dot, whereas individual values are indicated with smaller yellow dots.

**Figure 5.**
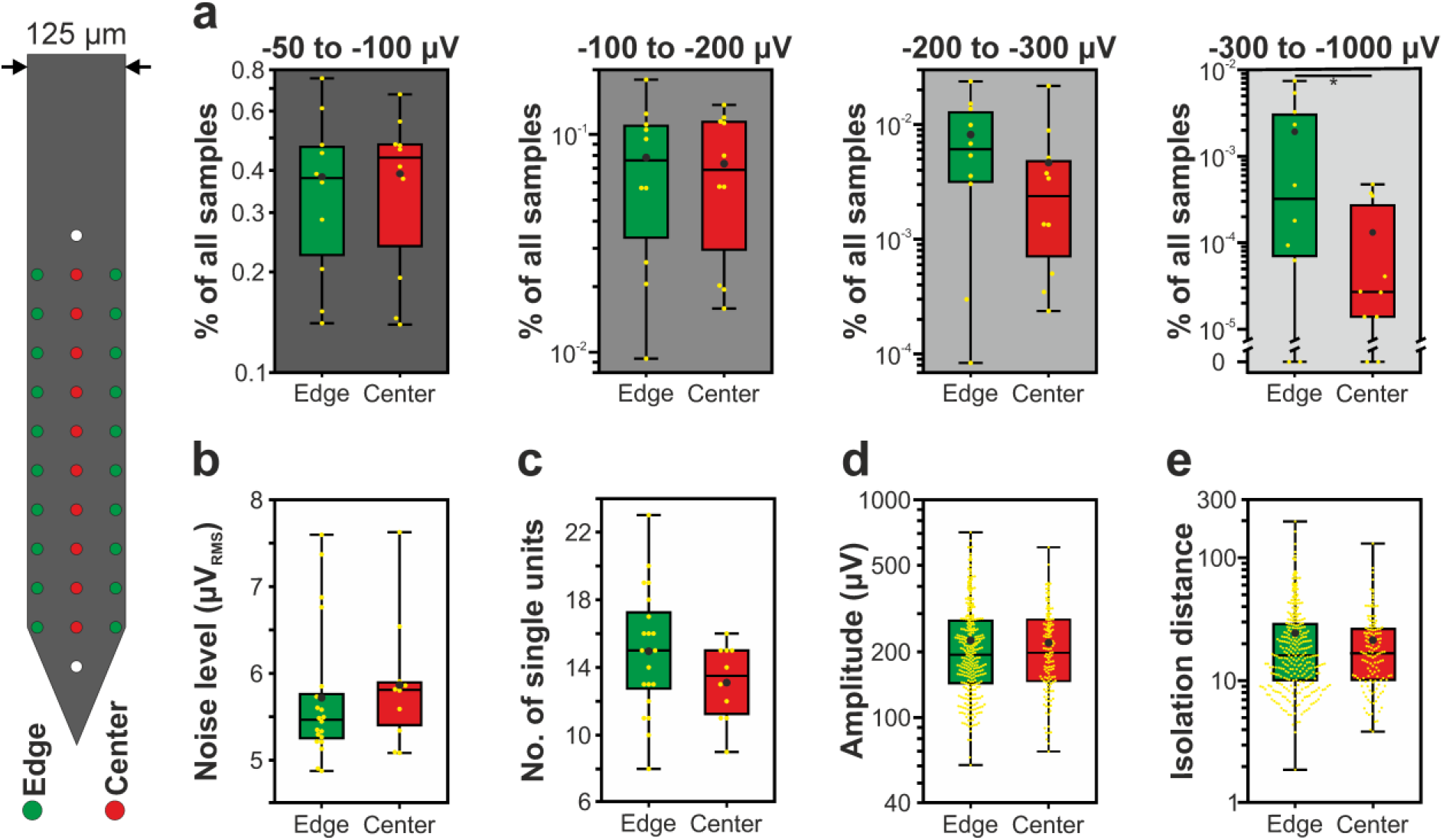
Boxplots showing the results of the 32-channel NeuroNexus silicon probe for edge (green) and center (red) sites. (a) Ratio of samples to the total number of samples for each of the four amplitude ranges (n = 10 recordings). (b) Estimated *in vivo* noise level. (c) Single unit yield (n = 430). (d) Peak-to-peak amplitude of the averaged single unit spike waveforms. (e) Isolation distance of the single unit clusters. The features of boxplots are demonstrated in figure 3(c). Note that most data are plotted on a logarithmic scale. * p < 0.05.

**Figure 6.**
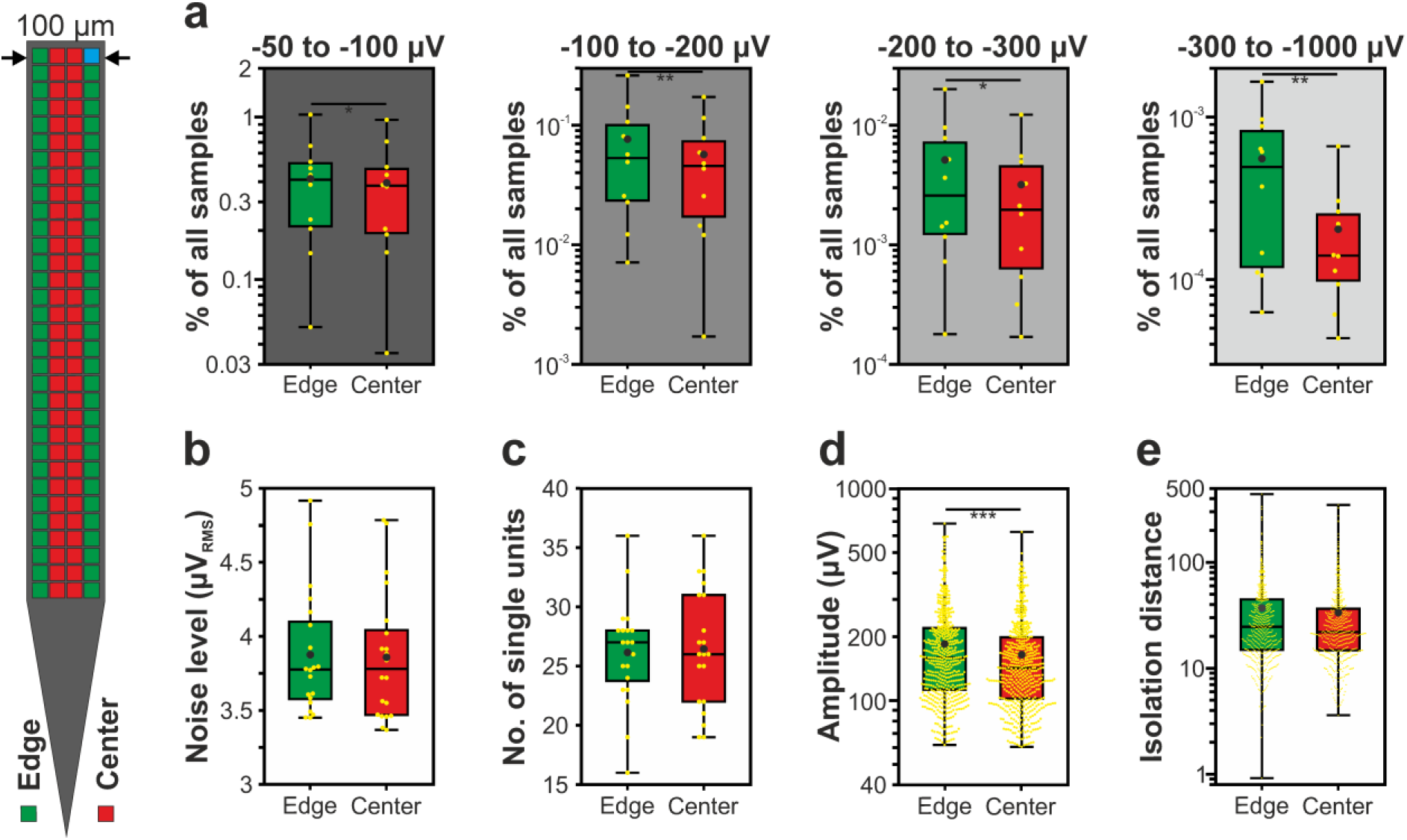
Boxplots showing the results of the 128-channel NeuroSeeker silicon probe for edge (green) and center (red) sites. (a) Ratio of samples to the total number of samples for each of the four amplitude ranges (n = 10 recordings). (b) Estimated *in vivo* noise level. (c) Single unit yield (n = 1052). (d) Peak-to-peak amplitude of the averaged single unit spike waveforms. (e) Isolation distance of the single unit clusters. The features of boxplots are demonstrated in figure 3(c). Note that most data are plotted on a logarithmic scale. * p < 0.05; ** p < 0.01; *** p < 0.001.

**Figure 7.**
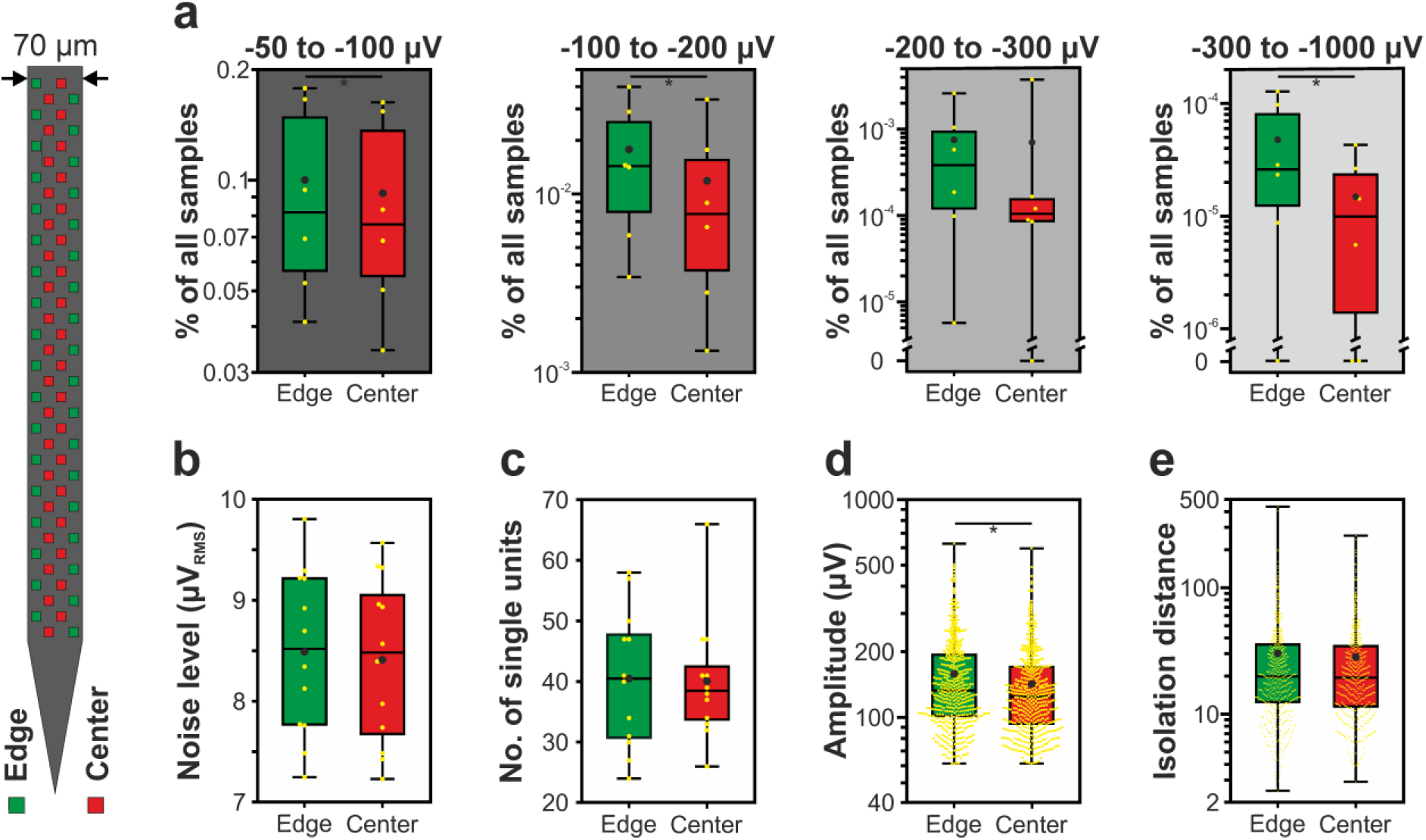
Boxplots showing the results of the 70-μm-wide Neuropixels silicon probe for edge (green) and center (red) sites. (a) Ratio of samples to the total number of samples for each of the four amplitude ranges (n = 6 recordings). (b) Estimated *in vivo* noise level. (c) Single unit yield (n = 967). (d) Peak-to-peak amplitude of the averaged single unit spike waveforms. (e) Isolation distance of the single unit clusters. The features of boxplots are demonstrated in figure 3(c). Note that most data are plotted on a logarithmic scale. * p < 0.05.

**Figure 8.**
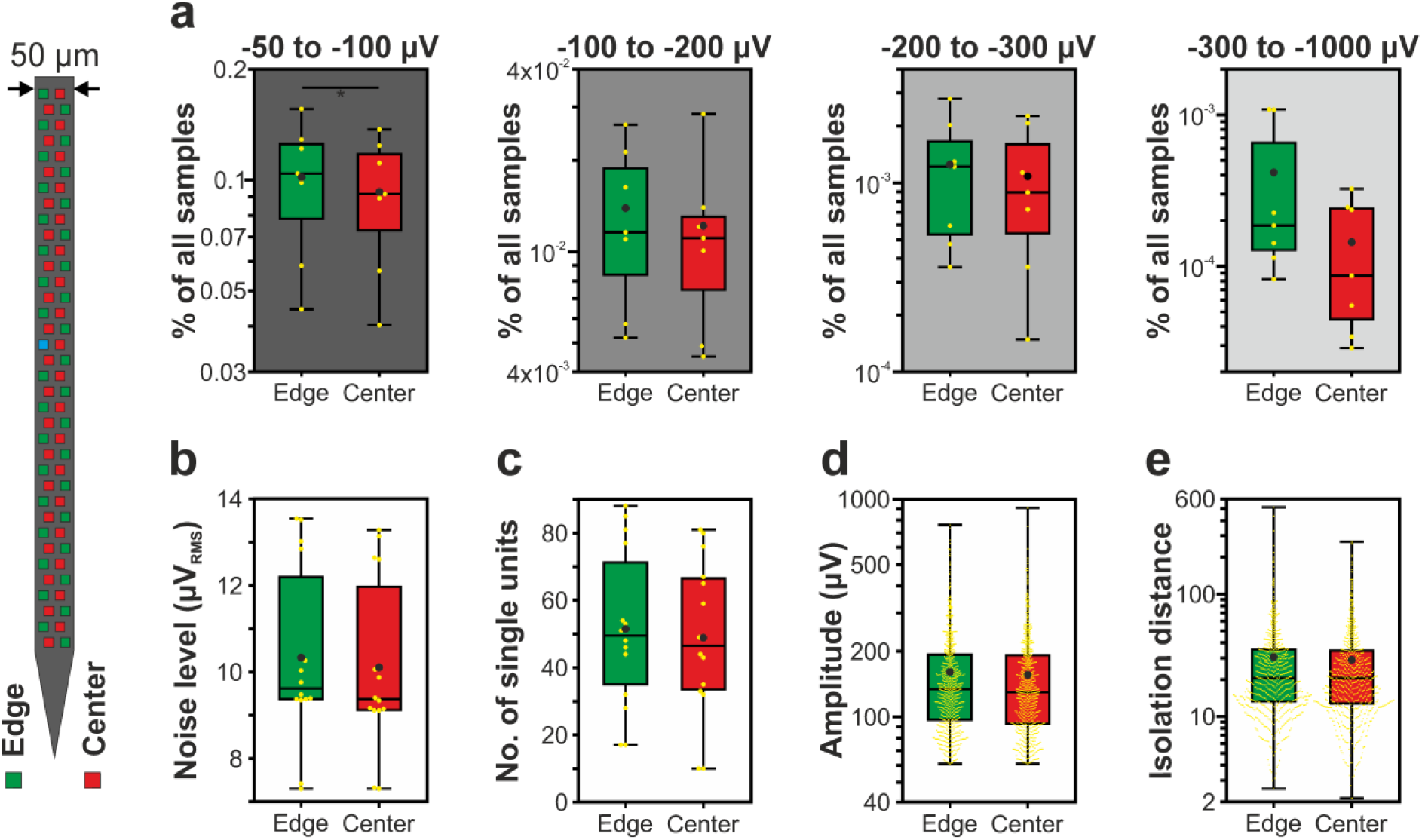
Boxplots showing the results of the 50-μm-wide Neuropixels silicon probe for edge (green) and center (red) sites. (a) Ratio of samples to the total number of samples for each of the four amplitude ranges (n = 7 recordings). (b) Estimated *in vivo* noise level. (c) Single unit yield (n = 1405). (d) Peak-to-peak amplitude of the averaged single unit spike waveforms. (e) Isolation distance of the single unit clusters. The features of boxplots are demonstrated in figure 3(c). Note that most data are plotted on a logarithmic scale. * p < 0.05.

**Figure 9.**
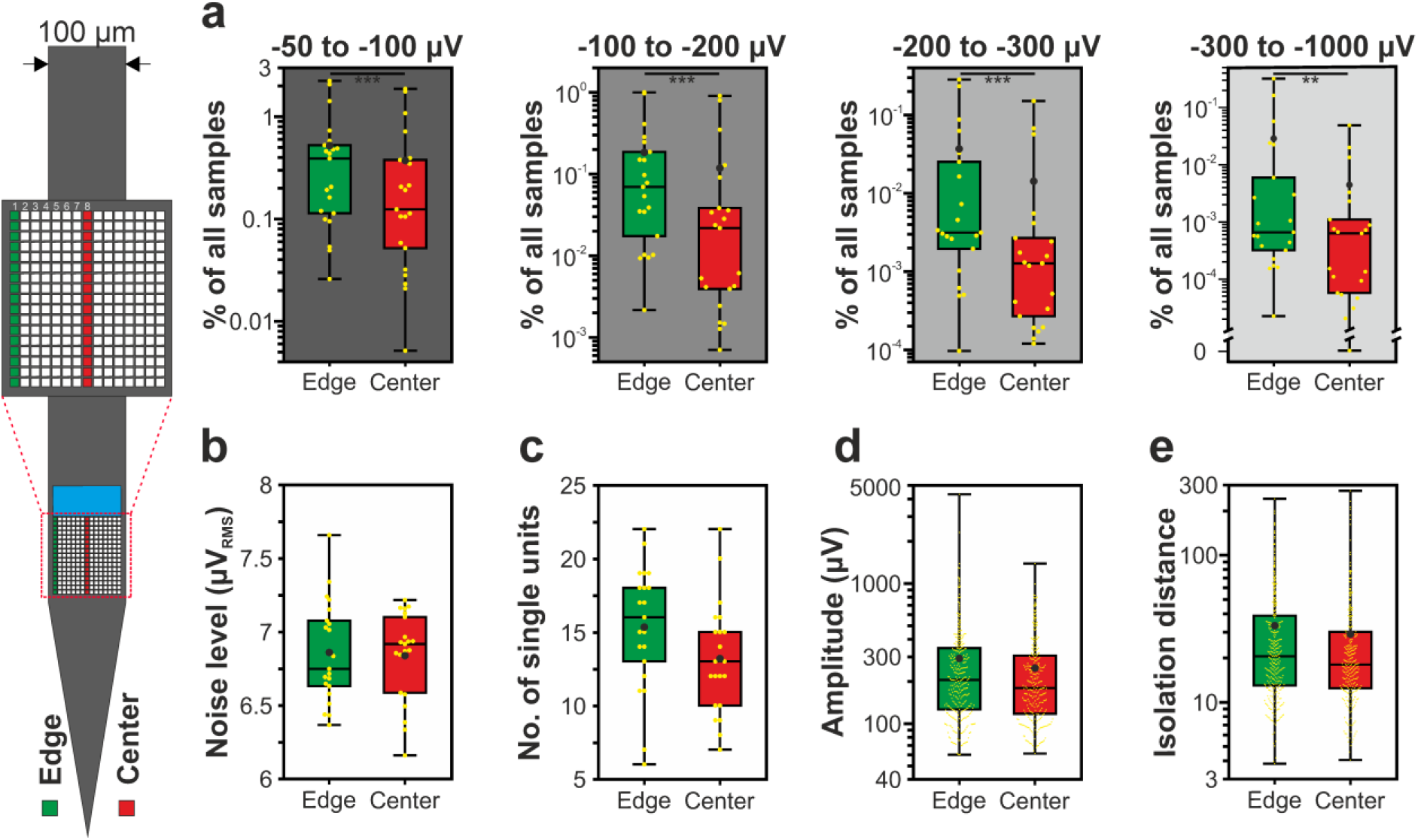
Boxplots showing the results of the 255-channel NeuroSeeker probe for edge (green) and center (red) sites. (a) Ratio of samples to the total number of samples for each of the four amplitude ranges (n = 21 recordings). (b) Estimated *in vivo* noise level. (c) Single unit yield (n = 599). (d) Peak-to-peak amplitude of the averaged single unit spike waveforms. (e) Isolation distance of the single unit clusters. The features of boxplots are demonstrated in figure 3(c). Note that most data are plotted on a logarithmic scale. ** p < 0.01; *** p < 0.001.

## 3. Results

To examine whether there is a difference in the recording performance of edge and center sites, we analyzed recordings obtained with commercially available and state-of-the-art high-density silicon probes with channel numbers ranging from 32 to 384 (NeuroNexus, Neuropixels and NeuroSeeker probes; figure 1; see section 2.1 for details). These single-shank planar probes contain recording sites both in the center and close to the edge of their silicon shank which makes them suitable to assess and compare the neural signal quality at these site positions. Furthermore, the different probe widths (from 50 μm to 125 μm) allow us to investigate width-dependent effects of the recording capability of edge and center sites. Quantitative details of *in vivo* experiments and cortical recordings are summarized in table 1.

To compare the signal quality provided by edge and center sites, we first separated channels of the spontaneous cortical recordings based on their site locations (figure 2; see section 2.5 for details), then extracted multiple features from the signals, focusing on the 500 – 5000 Hz frequency range corresponding to single unit activity. The amplitude of extracellular spikes quickly decays with distance [1, 60], that is, neurons located closer to recording sites will produce larger spikes which usually provides a better separability from spikes of other neurons. Thus, measures related to spike amplitude might be suitable to compare the quality of edge and center recordings. In addition, a higher proportion of high-amplitude spikes in the data might indirectly reflect a tissue damage of smaller extent, that is, the presence of more neurons which survived the probe insertion and are located close to a particular site [27]. Here, we examined the distribution of the amplitude values of the filtered potential in four non-overlapping negative amplitude ranges (figure 3; see section 2.6 for more details). To ascertain that differences in the amplitude distributions are not biased by differences in noise level between edge and center sites, we estimated the level of noise in the analyzed recordings based on the *in vivo* data (figure 4; see section 2.8 for more details). In addition, spike sorting was performed on the recordings to extract various single unit properties for comparison, such as the single unit yield, the amplitude of single unit spikes and the isolation distance (see section 2.7 for more details). The latter is a measure commonly used to determine the quality of single unit clusters [59], while the former are usual features used to assess the recording performance extracted under different conditions [27, 61]. Example recordings and single unit spike waveforms obtained with each probe at different site positions are shown in supplementary figures 2-6. Since we found no significant difference between the left and right sides of the probes, for each site position, we pooled the data corresponding to the two sides and did the analysis on the combined data (figures 5–9; tables 2–3). However, data of both sides can be found separately in the supplementary material (supplementary figures 7-10(a)-(e); supplementary tables 4-7).

**Table 2.** Mean ± standard deviation of the calculated features for all probe types (NN, NeuroNexus; NS, NeuroSeeker; NP, Neuropixels).

| | 32-channel NN probe | | 128-channel NS probe | | 70 $\mu\text{m}$ wide NP probe | | 50 $\mu\text{m}$ wide NP probe | | 255-channel NS probe | |
| --- | --- | --- | --- | --- | --- | --- | --- | --- | --- | --- |
|  | Edge | Center | Edge | Center | Edge | Center | Edge | Center | Edge | Center |
| –50 to –100 $\mu\text{V}$ (% of all samples) | $0.383 \pm 0.198$ | $0.391 \pm 0.179$ | $0.417 \pm 0.288$ | $0.393 \pm 0.279$ | $9.94 \times 10^{-2} \pm 5.81 \times 10^{-2}$ | $9.15 \times 10^{-2} \pm 5.35 \times 10^{-2}$ | $1.01 \times 10^{-1} \pm 3.90 \times 10^{-2}$ | $9.23 \times 10^{-2} \pm 3.48 \times 10^{-2}$ | $0.521 \pm 0.631$ | $0.373 \pm 0.547$ |
| –100 to –200 $\mu\text{V}$ (% of all samples) | $7.83 \times 10^{-2} \pm 5.37 \times 10^{-2}$ | $7.34 \times 10^{-2} \pm 4.59 \times 10^{-2}$ | $7.68 \times 10^{-2} \pm 7.83 \times 10^{-2}$ | $5.72 \times 10^{-2} \pm 5.32 \times 10^{-2}$ | $1.75 \times 10^{-2} \pm 1.38 \times 10^{-2}$ | $1.16 \times 10^{-2} \pm 1.20 \times 10^{-2}$ | $1.38 \times 10^{-2} \pm 7.78 \times 10^{-3}$ | $1.21 \times 10^{-2} \pm 7.97 \times 10^{-3}$ | $1.85 \times 10^{-1} \pm 2.89 \times 10^{-1}$ | $1.19 \times 10^{-1} \pm 2.57 \times 10^{-1}$ |
| –200 to –300 $\mu\text{V}$ (% of all samples) | $8.15 \times 10^{-3} \pm 7.51 \times 10^{-3}$ | $4.66 \times 10^{-3} \pm 6.55 \times 10^{-3}$ | $5.13 \times 10^{-3} \pm 6.13 \times 10^{-3}$ | $3.18 \times 10^{-3} \pm 3.70 \times 10^{-3}$ | $7.52 \times 10^{-4} \pm 9.82 \times 10^{-4}$ | $7.02 \times 10^{-4} \pm 1.49 \times 10^{-3}$ | $1.24 \times 10^{-3} \pm 8.85 \times 10^{-4}$ | $1.07 \times 10^{-3} \pm 8.01 \times 10^{-4}$ | $3.69 \times 10^{-2} \pm 7.74 \times 10^{-2}$ | $1.43 \times 10^{-2} \pm 3.63 \times 10^{-2}$ |
| –300 to –1000 $\mu\text{V}$ (% of all samples) | $1.92 \times 10^{-3} \pm 2.67 \times 10^{-3}$ | $1.32 \times 10^{-4} \pm 1.87 \times 10^{-4}$ | $5.54 \times 10^{-4} \pm 5.08 \times 10^{-4}$ | $2.04 \times 10^{-4} \pm 1.82 \times 10^{-4}$ | $4.51 \times 10^{-5} \pm 4.95 \times 10^{-5}$ | $1.41 \times 10^{-5} \pm 1.61 \times 10^{-5}$ | $4.10 \times 10^{-4} \pm 4.50 \times 10^{-4}$ | $1.42 \times 10^{-4} \pm 1.19 \times 10^{-4}$ | $2.87 \times 10^{-2} \pm 7.61 \times 10^{-2}$ | $4.45 \times 10^{-3} \pm 1.13 \times 10^{-2}$ |
| No. of single units | $14.95 \pm 3.73$ | $13.10 \pm 2.28$ | $26.15 \pm 4.44$ | $26.45 \pm 5.02$ | $40.50 \pm 11.49$ | $40.08 \pm 10.16$ | $51.50 \pm 23.85$ | $48.86 \pm 23.60$ | $15.33 \pm 4.23$ | $13.19 \pm 3.82$ |
| Amplitude of single units ($\mu\text{V}$) | $226.04 \pm 117.56$ | $220.11 \pm 97.39$ | $184.94 \pm 102.10$ | $164.65 \pm 85.37$ | $157.19 \pm 82.85$ | $140.93 \pm 70.14$ | $159.71 \pm 95.88$ | $154.97 \pm 92.76$ | $292.45 \pm 334.63$ | $249.39 \pm 191.66$ |
| Isolation distance | $24.38 \pm 23.86$ | $21.35 \pm 17.30$ | $37.08 \pm 42.14$ | $33.52 \pm 38.56$ | $30.25 \pm 37.02$ | $28.25 \pm 29.85$ | $30.71 \pm 37.73$ | $29.04 \pm 28.95$ | $33.20 \pm 34.66$ | $29.08 \pm 35.14$ |
| Noise level ($\mu\text{VRMS}$) | $5.72 \pm 0.79$ | $5.87 \pm 0.75$ | $3.88 \pm 0.42$ | $3.86 \pm 0.44$ | $8.49 \pm 0.82$ | $8.41 \pm 0.83$ | $10.34 \pm 2.09$ | $10.11 \pm 2.01$ | $6.88 \pm 0.45$ | $6.84 \pm 0.30$ |

**Table 3.**
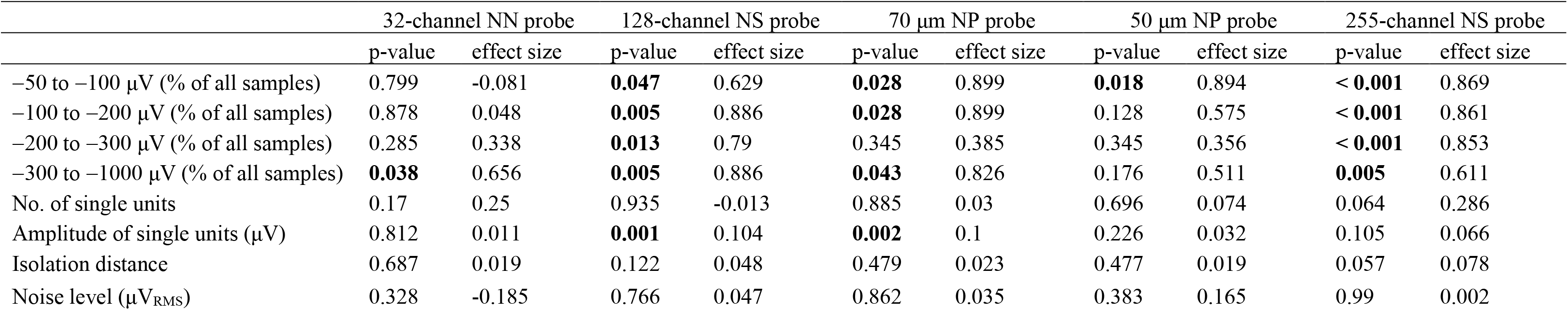
Summary of statistical analysis. Wilcoxon signed-rank test was applied for edge-center comparison in the four amplitude ranges, while Mann-Whitney U test was used in the case of the last four features. Significant (p < 0.05) values are indicated in bold (NN, NeuroNexus; NS, NeuroSeeker; NP, Neuropixels).

### 3.1 Comparison of the amplitude distribution of the filtered potential between different site positions

Boxplots corresponding to the analysis of the amplitude distribution are demonstrated in panel (a) of figures 5–9, while the mean values and the results of statistical analyses are summarized in tables 2 and 3, respectively. The noise level of recordings was low with a low variance across recordings, and there was no difference in the noise level of edge and center sites for either probe type (panel (b) of figures 5–9; tables 2 and 3). Examining the four amplitude ranges for each probe type, in almost all cases, edge sites provided a higher signal quality which was indicated by higher mean values of the examined features (95%, 19/20; table 2). Differences in the amplitude distribution between edge and center sites were significant in more than half of the cases with medium and large effect sizes (65%, 13/20; table 3), and all of these cases indicated an enhanced performance of edge sites. Edge sites significantly outperformed center sites for all probe types in at least one amplitude range, while in the case of the 128-channel and 255-channel probes, the difference was significant in all four amplitude intervals (figures 6(a) and 9(a); table 2). Based on the ratio of mean values of center sites to edge sites (table 2), as well as on the results of the statistical analysis (table 3), the difference between site positions was most remarkable in the range corresponding to the largest spikes (from −300 μV to −1000 μV; figure 10(a)). In this range, averaged across all probe types, mean values calculated for center sites were ~75% lower than mean values of edge sites (figure 10(a)). For the other three amplitude ranges, mean values of center sites were, on average, only ~22% lower compared to the mean values computed for edge sites (figure 10(a)). Similar results were obtained by investigating positive amplitude ranges (panel (f) of supplementary figures 7-10; supplementary figure 11(a); supplementary tables 4-8). That is, although the amount of positive values in the filtered recordings was usually much lower compared to the number of negative potential values (especially in higher amplitude ranges), mean values were in most cases higher for edge sites (95%, 19/20), and these differences were in about one third of the cases significant (30%, 6/20).

**Figure 10.**
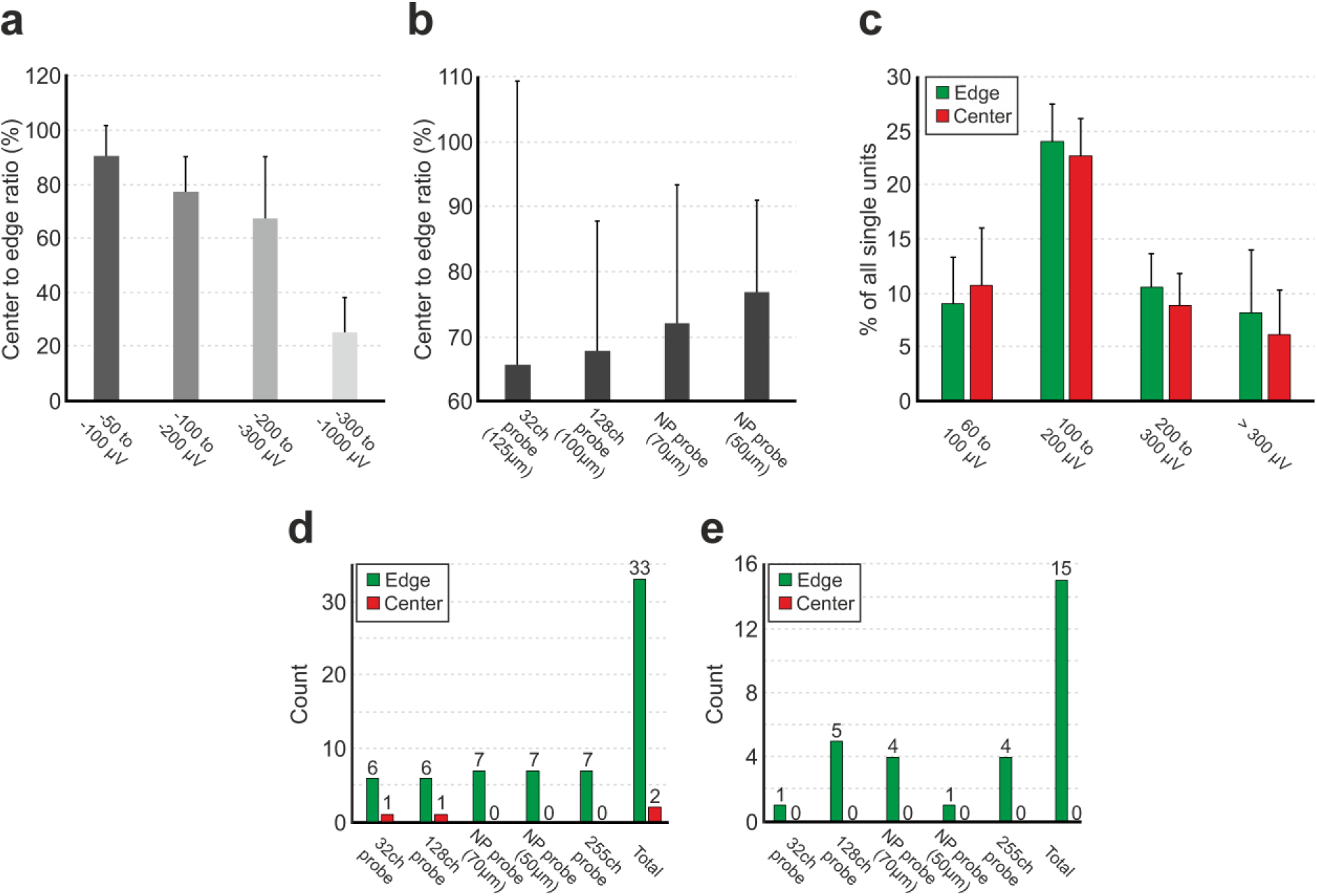
Summary of the recording performance comparison of edge and center sites. (a) Ratio of mean values of center sites to edge sites calculated for each amplitude range and averaged across probe types. (b) Ratio of mean values of center sites to edge sites calculated for each probe type (except the 255-channel probe) and averaged across all amplitude ranges. Probes are ordered by shank width. (c) Ratio of single units to the total number of units calculated for each site position in four ranges of the peak-to-peak amplitude of spike waveforms. Results are averaged across probe types. (d) Quantity of higher mean values of the examined seven features (four amplitude ranges, single unit yield, unit amplitude and isolation distance) compared between edge and center sites, shown for all five probe types. (e) Number of examined features where a statistically significant difference was found between edge and center sites, shown for all five probes types.

To examine whether there might be shank width-dependent differences in the recording performance between edge and center sites, for each probe type, we averaged the ratio of center-to-edge mean values across all amplitude ranges. Results of the 255-channel probe were not used in this analysis, since this probe type has the same width as the 128-channel probe and a significantly different site configuration. Our results show, that with decreasing shank width, a slightly increasing trend in the ratio of mean values of center sites to edge sites can be observed (figure 10(b)). This suggests that the performance advantage of edge sites becomes smaller for narrower probes.

By investigating recordings obtained with the 255-channel probe providing a superior spatial resolution (figure 9), we can perform a finer and more detailed analysis to compare the recording performance of sites located at different distances from the edge of the silicon shank. Therefore, we analyzed the amplitude distribution of recordings obtained at eight adjacent columns of recording sites (17 sites in each column) located on the left side of probe (supplementary figure 11(b); supplementary table 9). For each amplitude range, the highest mean value was provided by the first column of sites (located at the edge), whereas the lowest values were measured at the recording sites located in the center of the shank. Values in each amplitude range showed a slightly decreasing trend from edge sites towards the center of the silicon shank.

To further investigate the robustness of our results, we analyzed a public dataset (n = 7 cortical recordings) obtained with the same type of 255-channel probe as used in this study (supplementary figure 12(a); supplementary table 10). Mean values of edge sites were higher in three amplitude ranges compared to values of center sites (with one significantly higher), while in the range containing the largest spikes, center sites performed slightly better. However, based on the mean values of the amplitude distribution and spike sorting results, these recordings had a lower quality compared to our recordings (figure 9 vs. supplementary figure 12), probably due to differences in the anesthesia, insertion conditions, or targeted brain region. Nevertheless, the overall signal quality was still somewhat better at edge sites than at center sites.

The relatively large effect sizes obtained here (table 3) might also be the results of low sample sizes (~10 recordings/probe). To examine the influence of sample size on the effect size, we analyzed a larger dataset (n = 186 cortical recordings) acquired with the 128-channel NeuroSeeker probe. Although the effect sizes decreased (from 0.80 ± 0.12 to 0.58 ± 0.08; averaging across the four amplitude ranges), these still indicated a medium effect, and the differences between edge and center sites were also highly significant (p < 0.001 for all amplitude ranges; supplementary figure 13; supplementary tables 11-12).

The neocortex has a special structure with multiple layers, areas, columns and various cell types [62]. To examine whether there might by brain area-dependent differences in the recording performance of edge and center sites, we analyzed recordings (n = 9) from the somatosensory thalamus obtained with the 128-channel probe (supplementary figure 14; supplementary tables 13-14). Again, edge sites provided better signal quality compared to center sites, including all investigated amplitude ranges, although the difference in recording performance was smaller. Significant differences were only found between site positions in the two amplitude ranges containing the largest thalamic spikes (supplementary figure 14(a)).

It would be interesting to examine whether there are differences in the recording performance between sites located at different longitudinal positions of the silicon shank. To investigate that, we analyzed the same cortical (n = 10) and thalamic (n = 9) recordings obtained with the 128-channel silicon probes (supplementary figure 15; supplementary tables 15-16). However, in this case, four site groups (32 channels in each group) were created based on their vertical locations (supplementary figure 15(a)). After that, we calculated the ratio of amplitude values in the four amplitude ranges (supplementary figure 15(b) and (c)). For recordings from both brain structures, the results showed that the site group located in the lower middle of the shank (second group of 32-channels, calculated from the bottom) provided the best recording performance, in all amplitude ranges (supplementary figure 15(b) and (c); supplementary tables 15-16). However, the results might be slightly biased in the case of cortical recordings, since the intensity of spiking activity significantly varies across cortical layers in ketamine/xylazine anesthetized rats (supplementary figure 15(d); [57]). Unit activity was found to be the strongest in the lower part of layer V which has a thickness about 300 μm, and was weaker in upper and lower layers [57]. Because recordings with the 128-channel probe were done in multiple cortical layers simultaneously (usually layers IV-VI), the layer-dependent intensity of spiking activity might affect our results obtained here. However, results obtained with thalamic recordings might provide a more accurate comparison because the recorded spiking activity was more uniform at different depths compared to cortical activity (supplementary figure 15(d)). Although small differences in the structure of the thalamic area examined or depth-dependent differences in thalamic activity (e.g. by recording simultaneously from multiple thalamic nuclei) may still slightly decrease the reliability of these findings.

In summary, edge sites provided a higher performance compared to center sites in all amplitude ranges and for all probe types. Such differences in signal quality could be observed in different brain areas, and also at different longitudinal positions of the recording sites. The better performance of the edge sites decreased with decreasing shank width.

### 3.2 Comparison of single unit properties between edge and center sites

Boxplots corresponding to the analysis of single units are demonstrated in panels (c)-(e) of figures 5–9, while mean values and the results of statistical analyses are summarized in tables 2 and 3, respectively. In almost all cases, edge sites provided a higher signal quality which was indicated by higher mean values of the single unit yield, spike amplitude and isolation distance (~93%, 14/15; table 2). The average number of well-separated single units recorded at a particular brain location ranged from ~15 (32-channel NeuroNexus and 255-channel NeuroSeeker probes) to ~50 (50-μm-wide Neuropixels probe). The difference in the single unit yield was small and not significant between site positions, usually only a few more units could be separated on edge sites. However, the spike amplitude of single units obtained with the 128-channel NeuroSeeker probe and the wider Neuropixels probe at edge sites were significantly higher. Overall, more single units having spike waveforms with peak-to-peak amplitudes over 100 μV were detected on edge sites (figure 10(c)). This suggests that either more neurons were located closer to edge sites than to center sites (e.g. due to tissue compression caused by the probe) or more neurons survived the mechanical trauma of probe insertion in the vicinity of edge sites.

## 4. Discussion

Our results, based on quantitative analysis of high-density neural recordings, showed that, for all probe types examined, the recording performance of edge sites was notably better than that of center sites. Examined features of the filtered signal containing spiking activity had in most cases higher mean values for edge sites (~94%, 33/35; figure 10(d)). Differences between the two site positions were also significant in several cases (~45%, 15/35; figure 10(e)), and these differences indicated in every case a better signal quality recorded on edge sites. These differences were most remarkable in the amplitude range corresponding to large (with < −300 μV peak) spikes (figure 10(a)). Furthermore, we have found that the shank width of the probe might also affect the difference in recording performance: edge sites lose their advantage with narrower shanks (figure 10(b)). Although the single unit yield was not significantly higher between the site positions, the amplitude of the unit spike waveforms was usually larger, and the quality of unit clusters was better for edge sites (figure 10(c)). The small difference in the single unit yield and the small effect sizes found in the case of single unit features might suggest only a moderate practical applicability of our findings; however, the significantly larger spikes recorded at edge sites may notably improve the separability of single unit clusters, and thus the accuracy and reliability of spike sorting results.

### 4.1 Comparison with previous work

Out of three studies investigating the recording performance of microelectrodes located on the edge and the center of silicon shanks [34–36], the results of two studies are in agreement with our findings, that is, placing recording sites on the edge improves the quality of the recorded neural signal. Lee and colleagues chronically implanted custom-designed 16-channel silicon probes with two different widths in the motor cortex of rats to track their recording performance for several weeks [35]. Seven and eight quadratic recording sites (30 μm × 30 μm) were placed in the center and at the edge of the probe shank, respectively. Compared to center sites, the chronic recording capability of edge sites of the wider probe was significantly better for several weeks in terms of the ratio of sites containing spiking activity as well as the signal-to-noise ratio of separated single units. Although the results were not significant, edge sites of the planar silicon probe with the narrower shank (132 μm) still performed better than center sites. Our comprehensive study extended these results by analyzing data obtained with probes having narrower shanks (from 50 μm to 125 μm). Interestingly, edge sites still provided better recordings even with the narrowest device, although the advantage of edge sites decreased notably with decreasing shank widths.

The custom-designed probes developed by Seymour and colleagues [36] had microelectrodes fabricated in the lateral wall of the silicon shank (edge sites) in combination with recording sites placed on the front and back side of the shank located further from the shank edge. They could separate more single units in recordings obtained with edge sites and the measured spike amplitudes were also higher. Although we found no difference in the single unit yield between edge and center sites, edge sites on the investigated high-density probes are located not exactly on the edge but a few micrometers towards the center. In contrast, the special edge design used in [36] might provide a better accessibility to neurons and a wider viewing angle to detect their action potentials due to a decreased shielding effect of the silicon shank [63, 64].

In the work of Scott et al. [34], an 85 μm wide single-shank silicon probe with 64 recording sites arranged in three columns (two on each side and one in the center of the shank) was used to record spiking activity from the hippocampus of mice. They found no difference in the signal quality between edge and center sites, as well as between spike amplitudes measured on sites located at different longitudinal positions of the shank. The contradiction between their results and our findings might be caused by several factors such as the difference in animal species, the examined brain area or the used methodology.

### 4.2 Limitations of the study

As described above, recording sites classified as edge sites in this study are located slightly further (~5-6 μm) from the edge of the silicon shank. Thus, our results obtained with edge sites might slightly underestimate the real recording performance of microelectrodes placed exactly at the edge of the shank, for example, in the case of special edge probe designs [36, 41, 65] which provide an even larger field of view for signal detection.

The focus of this study was on the analysis of cortical recordings. However, as demonstrated by comparing the performance of edge and center sites between thalamic and cortical recordings (with a larger difference in the signal quality in the neocortex), the performance advantage of edge sites might also depend on the structure, cellular density and composition of the examined brain region. This theory is further supported by the findings of the study of Scott and colleagues, where no difference was found in the recording capabilities of edge and center sites in another brain region, namely in the hippocampus of mice [34]. In contrast, in the case of the two studies which have found a difference in the signal quality between edge and center sites, the probes were implanted into the motor cortex of rats [35, 36]. Therefore, testing the recording performance of edge and center sites in other brain areas as well as in other species might further our knowledge of optimal site placement on silicon probes.

Other factors might also influence the quality of recordings, and thus might affect the performance difference between edge and center sites. For instance, a computational modeling study demonstrated that both edema and glial encapsulation can have a significant impact on the amplitude of the recorded spikes, with a decrease of amplitude in the former case and an increase in the latter case [63]. Thus, a localized edema developed due to a minor tissue damage near the edge of the shank might diminish the performance advantage of affected edge sites. Moreover, a glial scar formed around chronically implanted silicon probes might be more pronounced close to the sharp edges of the probe where small micromotions of the implant mechanically insult the tissue [66, 67]. This difference in glial density or thickness might influence the signal quality of recording sites, especially those located at the edges. Although we only analyzed recordings of acute experiments with a relatively short duration (~30 min), results of the work of Lee et al. [35] suggest that edge sites might keep their performance advantage over center sites also over longer timescales, even for several weeks.

The speed of probe insertion may also significantly affect recording quality [27]. Analyzing the dataset of our previous study [27], we found that, in acute recordings, the performance difference between edge and center sites was larger when slower speeds (< 0.1 mm/sec) were used for insertion (data not shown). As the speed of insertion affects the extent of neuronal loss around the probe [27], this suggests that using a higher insertion speed will result in more severe tissue damage close to the edges of the probe.

### 4.3 Possible interpretations of the enhanced signal quality provided by edge sites

Based on our current knowledge, we can only speculate on the causes behind the observed differences in recording performance of edge and center sites. One plausible explanation might be that edge sites have a better “visibility” compared to center sites, that is, they are less affected by the shielding or shadowing effect of the silicon substrate [36, 63]. Therefore, microelectrodes located closer to the edge of the shank should detect the action potentials of more neurons. However, we did not find a higher single unit yield for edge sites. The main difference in the recording performance between the two site positions was in the amplitude of spikes, mostly in high amplitude ranges corresponding to the largest spikes. Because the spike amplitude rapidly attenuates with increasing distance from the soma of the neuron [1, 60], higher spike amplitudes on edge sites may suggest that several neurons are located closer to edge sites than to center sites. The reason behind this asymmetry in the distance of neurons might be that more neurons survive the implantation of the probe which are close to the edge of the shank compared to cells located close to the center. However, this scenario should probably also result in a higher unit yield for edge sites. Another possible explanation might be that, since the width of these probes is larger than their thickness, the tissue compression caused by the probe is asymmetric and is higher along the lateral axis than along the axis corresponding to the front and backside of the probe. Thus, neurons might be slightly more compressed near edge sites and pushed closer to them. Therefore, using a probe shank with a smaller cross-section should result in the decrease of performance difference between the two site positions, as the degree of tissue compression will be smaller. We could observe this decrease in performance: although still present, the advantage of edge site over center sites slightly decreased with decreasing shank width.

It is also important to mention, that the shielding effect of the silicon shank may also affect the recorded spike amplitudes [36, 46, 63]. One computational modeling study showed that when a modeled probe shank is placed before the model neuron, the recorded spike amplitudes are almost two times higher compared to the amplitude of simulated spikes obtained without the presence of the probe shank [63]. In a recent modeling study, using mesh models of NeuroNexus and Neuropixels probes, Buccino and colleagues demonstrated similar findings, that is, the use of silicon probes significantly amplified the recorded potentials [46]. They have also found that almost two times higher action potential amplitudes can be detected when the probe shank is present. This difference in the spike amplitudes decreased when the model neuron was shifted from the center of the shank laterally to the edge. For instance, in the case a 50 μm lateral shift, the simulated spike amplitudes were only 40% higher with the probe shank included in the simulations. The authors argue that this bias might results in more neurons found in the probe center than at the edges. If these differences in the recorded spike amplitudes are similar in *in vivo* recordings, then, based on our results, the recording performance at edge sites should be even better than our analysis showed.

### 4.3 Recommendations for recording site placement in novel electrode designs

Silicon microprobes are extensively used in a many electrophysiology labs. Further advancements in probe design and in silicon microfabrication technology will soon make it possible for research groups to design they own recording devices with features customized to the actual research task. This process is further facilitated by recently shared open source probe designs and open source hardware for electrophysiological experiments [37, 40]. Our results might help engineers and scientists working in the field of neuroscience to determine the optimal placement of microelectrodes on the shank of planar silicon probes. For instance, placing recording sites on the edge of the shank of passive probes which have a limited site number may significantly enhance their recording performance. This can save time needed to perform experiments and also reduce the costs of these studies. From another aspect, our findings suggest that in the case of high-density probes with a small shank cross-section (e.g. the Neuropixels probe), where the entire shank is covered with recording sites, both edge and center sites will sample the neuronal activity with similar quality and recordings will not be biased towards either of these site positions.

## Acknowledgements

The research leading to these results has received funding from the Hungarian Brain Research Program Grant (Grant No. 2017-1.2.1-NKP-2017-00002) and from the European Union’s Seventh Framework Program (FP7/2007-2013) under grant agreement n°600925 (NeuroSeeker). The research within project No. VEKOP-2.3.2-16-2017-00013 by I. Ulbert was supported by the European Union and the State of Hungary, co-financed by the European Regional Development Fund. R. Fiáth is thankful to the Hungarian National Research, Development and Innovation Office (PD124175).

## References

[1] Buzsaki G 2004 Large-scale recording of neuronal ensembles Nat. Neurosci. 7 446–51

[2] Buzsaki G, Stark E, Berenyi A, Khodagholy D, Kipke D R, Yoon E and Wise K D 2015 Tools for Probing Local Circuits: High-Density Silicon Probes Combined with Optogenetics Neuron 86 92–105

[3] Jun J J, Steinmetz N A, Siegle J H, Denman D J, Bauza M, Barbarits B, Lee A K, Anastassiou C A, Andrei A, Aydin C, Barbic M, Blanche T J, Bonin V, Couto J, Dutta B, Gratiy S L, Gutnisky D A, Hausser M, Karsh B, Ledochowitsch P, Lopez C M, Mitelut C, Musa S, Okun M, Pachitariu M, Putzeys J, Rich P D, Rossant C, Sun W L, Svoboda K, Carandini M, Harris K D, Koch C, O’Keefe J and Harris T D 2017 Fully integrated silicon probes for high-density recording of neural activity Nature 551 232–6

[4] Raducanu B C, Yazicioglu R F, Lopez C M, Ballini M, Putzeys J, Wang S W, Andrei A, Rochus V, Welkenhuysen M, van Helleputte N, Musa S, Puers R, Kloosterman F, Van Hoof C, Fiath R, Ulbert I and Mitra S 2017 Time Multiplexed Active Neural Probe with 1356 Parallel Recording Sites Sensors 17 2388

[5] Steinmetz N A, Koch C, Harris K D and Carandini M 2018 Challenges and opportunities for large-scale electrophysiology with Neuropixels probes Curr. Opin. Neurobiol. 50 92–100

[6] Steinmetz N A, Zatka-Haas P, Carandini M and Harris K D 2019 Distributed coding of choice, action and engagement across the mouse brain Nature 576 266–73

[7] Harris K D, Quiroga R Q, Freeman J and Smith S L 2016 Improving data quality in neuronal population recordings Nat. Neurosci. 19 1165–74

[8] Marton G, Baracskay P, Cseri B, Plosz B, Juhasz G, Fekete Z and Pongracz A 2016 A silicon-based microelectrode array with a microdrive for monitoring brainstem regions of freely moving rats Journal of neural engineering 13 026025

[9] Stringer C, Pachitariu M, Steinmetz N, Carandini M and Harris K D 2019 High-dimensional geometry of population responses in visual cortex Nature 571 361–5

[10] Stringer C, Pachitariu M, Steinmetz N, Reddy C B, Carandini M and Harris K D 2019 Spontaneous behaviors drive multidimensional, brainwide activity Science 364 255

[11] Yang W and Yuste R 2017 In vivo imaging of neural activity Nat. Methods 14 349–59

[12] Lin M Z and Schnitzer M J 2016 Genetically encoded indicators of neuronal activity Nat. Neurosci. 19 1142–53

[13] Weisenburger S and Vaziri A 2018 A Guide to Emerging Technologies for Large-Scale and Whole-Brain Optical Imaging of Neuronal Activity Annu. Rev. Neurosci. 41 431–52

[14] Fiath R, Raducanu B C, Musa S, Andrei A, Lopez C M, van Hoof C, Ruther P, Aarts A, Horvath D and Ulbert I 2018 A silicon-based neural probe with densely-packed low-impedance titanium nitride microelectrodes for ultrahigh-resolution in vivo recordings Biosens. Bioelectron. 106 86–92

[15] Putzeys J, Raducanu B C, Carton A, De Ceulaer J, Karsh B, Siegle J H, Van Helleputte N, Harris T D, Dutta B, Musa S and Lopez C M 2019 Neuropixels Data-Acquisition System: A Scalable Platform for Parallel Recording of 10,000+ Electrophysiological Signals IEEE transactions on biomedical circuits and systems 13 1635–44

[16] Wang S, Garakoui S K, Chun H, Gomez Salinas D, Sijbers W, Putzeys J, Martens E, Craninckx J, Van Helleputte N and Mora Lopez C 2019 A Compact Quad-Shank CMOS Neural Probe with 5,120 Addressable Recording Sites and 384 Fully Differential Parallel Channels IEEE transactions on biomedical circuits and systems 13 1625–34

[17] Angotzi G N, Boi F, Lecomte A, Miele E, Malerba M, Zucca S, Casile A and Berdondini L 2019 SiNAPS: An implantable active pixel sensor CMOS-probe for simultaneous large-scale neural recordings Biosens. Bioelectron. 126 355–64

[18] Herbawi A S, Christ O, Kiessner L, Mottaghi S, Hofmann U G, Paul O and Ruther P 2018 CMOS Neural Probe With 1600 Close-Packed Recording Sites and 32 Analog Output Channels Journal of Microelectromechanical Systems 27 1023–34

[19] De Dorigo D, Moranz C, Graf H, Marx M, Wendler D, Shui B Y, Herbawi A S, Kuhl M, Ruther P, Paul O and Manoli Y 2018 Fully Immersible Subcortical Neural Probes With Modular Architecture and a Delta-Sigma ADC Integrated Under Each Electrode for Parallel Readout of 144 Recording Sites IEEE JSolid-St Circ 53 3111–25

[20] Ruther P and Paul O 2015 New approaches for CMOS-based devices for large-scale neural recording Curr. Opin. Neurobiol. 32 31–7

[21] Karumbaiah L, Saxena T, Carlson D, Patil K, Patkar R, Gaupp E A, Betancur M, Stanley G B, Carin L and Bellamkonda R V 2013 Relationship between intracortical electrode design and chronic recording function Biomaterials 34 8061–74

[22] Thelin J, Jorntell H, Psouni E, Garwicz M, Schouenborg J, Danielsen N and Linsmeier C E 2011 Implant size and fixation mode strongly influence tissue reactions in the CNS PLoS One 6 e16267

[23] Szarowski D H, Andersen M D, Retterer S, Spence A J, Isaacson M, Craighead H G, Turner J N and Shain W 2003 Brain responses to micro-machined silicon devices Brain Res. 983 23–35

[24] Edell D J, Toi V V, Mcneil V M and Clark L D 1992 Factors Influencing the Biocompatibility of Insertable Silicon Microshafts in Cerebral-Cortex IEEE Trans. Biomed. Eng. 39 635–43

[25] Ersen A, Elkabes S, Freedman D S and Sahin M 2015 Chronic tissue response to untethered microelectrode implants in the rat brain and spinal cord Journal of neural engineering 12 016019

[26] Biran R, Martin D C and Tresco P A 2007 The brain tissue response to implanted silicon microelectrode arrays is increased when the device is tethered to the skull J. Biomed. Mater. Res. A 82 169–78

[27] Fiath R, Marton A L, Matyas F, Pinke D, Marton G, Toth K and Ulbert I 2019 Slow insertion of silicon probes improves the quality of acute neuronal recordings Sci. Rep. 9 111

[28] Bjornsson C S, Oh S J, Al-Kofahi Y A, Lim Y J, Smith K L, Turner J N, De S, Roysam B, Shain W and Kim S J 2006 Effects of insertion conditions on tissue strain and vascular damage during neuroprosthetic device insertion Journal of neural engineering 3 196–207

[29] Welkenhuysen M, Andrei A, Ameye L, Eberle W and Nuttin B 2011 Effect of insertion speed on tissue response and insertion mechanics of a chronically implanted silicon-based neural probe IEEE Trans. Biomed. Eng. 58 3250–9

[30] Neto J P, Baiao P, Lopes G, Frazao J, Nogueira J, Fortunato E, Barquinha P and Kampff A R 2018 Does Impedance Matter When Recording Spikes With Polytrodes? Front. Neurosci. 12 715

[31] Viswam V, Obien M E J, Franke F, Frey U and Hierlemann A 2019 Optimal Electrode Size for Multi-Scale Extracellular-Potential Recording From Neuronal Assemblies Front. Neurosci. 13 385

[32] Cogan S F 2008 Neural stimulation and recording electrodes Annu. Rev. Biomed. Eng. 10 275–309

[33] Negi S, Bhandari R and Solzbacher F 2012 Morphology and Electrochemical Properties of Activated and Sputtered Iridium Oxide Films for Functional Electrostimulation Journal of Sensor Technology 02 138–47

[34] Scott K M, Du J, Lester H A and Masmanidis S C 2012 Variability of acute extracellular action potential measurements with multisite silicon probes J. Neurosci. Methods 211 22–30

[35] Lee H C, Gaire J, Roysam B and Otto K J 2018 Placing Sites on the Edge of Planar Silicon Microelectrodes Enhances Chronic Recording Functionality IEEE Trans. Biomed. Eng. 65 1245–55

[36] Seymour J P, Langhals N B, Anderson D J and Kipke D R 2011 Novel multi-sided, microelectrode arrays for implantable neural applications Biomed. Microdevices 13 441–51

[37] Siegle J H, Cuevas Lopez A, Patel Y, Abramov K, Ohayon S and Voigts J 2017 Open Ephys: An open-source, plugin-based platform for multichannel electrophysiology Journal of neural engineering 14 045003

[38] Solari N, Sviatkó K, Laszlovszky T, Hegedüs P and Hangya B 2018 Open Source Tools for Temporally Controlled Rodent Behavior Suitable for Electrophysiology and Optogenetic Manipulations Front. Syst. Neurosci. 12 18

[39] Nasiotis K, Cousineau M, Tadel F, Peyrache A, Leahy R M, Pack C C and Baillet S 2019 Integrated open-source software for multiscale electrophysiology Sci Data 6 231

[40] Yang L, Lee K, Villagracia J and Masmanidis S C 2020 Open source silicon microprobes for high throughput neural recording Journal of neural engineering 17 016036

[41] Ulyanova A V, Cottone C, Adam C D, Gagnon K G, Cullen D K, Holtzman T, Jamieson B G, Koch P F, Chen H I, Johnson V E and Wolf J A 2019 Multichannel Silicon Probes for Awake Hippocampal Recordings in Large Animals Front. Neurosci. 13 397

[42] Dimitriadis G, Neto J P, Aarts A, Alexandru A, Ballini M, Battaglia F, Calcaterra L, David F, Fiath R, Frazao J, Geerts J P, Gentet L J, Helleputte N V, Holzhammer T, Hoof C v, Horvath D, Lopes G, Lopez C M, Maris E, Marques-Smith A, Marton G, McNaughton B L, Meszena D, Mitra S, Musa S, Neves H, Nogueira J, Orban G A, Pothof F, Putzeys J, Raducanu B, Ruther P, Schroeder T, Singer W, Tiesinga P, Ulbert I, Wang S, Welkenhuysen M and Kampff A R 2018 Why not record from every channel with a CMOS scanning probe? bioRxiv 275818

[43] Neto J P, Lopes G, Frazao J, Nogueira J, Lacerda P, Baiao P, Aarts A, Andrei A, Musa S, Fortunato E, Barquinha P and Kampff A R 2016 Validating silicon polytrodes with paired juxtacellular recordings: method and dataset J. Neurophysiol. 116 892–903

[44] English D F, McKenzie S, Evans T, Kim K, Yoon E and Buzsaki G 2017 Pyramidal Cell-Interneuron Circuit Architecture and Dynamics in Hippocampal Networks Neuron 96 505–20 e7

[45] Laboy-Juarez K J, Ahn S and Feldman D E 2019 A normalized template matching method for improving spike detection in extracellular voltage recordings Sci. Rep. 9 12087

[46] Buccino A P, Kuchta M, Jaeger K H, Ness T V, Berthet P, Mardal K A, Cauwenberghs G and Tveito A 2019 How does the presence of neural probes affect extracellular potentials? Journal of neural engineering 16 026030

[47] Klee J L, Kiliaan A J, Lipponen A and Battaglia F P 2019 Reduced firing rates of pyramidal cells in frontal cortex of APP/PS1 can be restored by acute treatment with levetiracetam bioRxiv 739912

[48] Klein L, Pothof F, Raducanu B C, Klon-Lipok J, Shapcott K A, Musa S, Andrei A, Aarts A A, Paul O, Singer W and Ruther P 2020 High-density electrophysiological recordings in macaque using a chronically implanted 128-channel passive silicon probe Journal of neural engineering 17 026036

[49] Fiath R, Raducanu B C, Musa S, Andrei A, Lopez C M, Welkenhuysen M, Ruther P, Aarts A and Ulbert I 2019 Fine-scale mapping of cortical laminar activity during sleep slow oscillations using high-density linear silicon probes J. Neurosci. Methods 316 58–70

[50] Marques-Smith A, Neto J P, Lopes G, Nogueira J, Calcaterra L, Frazão J, Kim D, Phillips M G, Dimitriadis G and Kampff A 2018 Recording from the same neuron with high-density CMOS probes and patch-clamp: a ground-truth dataset and an experiment in collaboration bioRxiv 370080

[51] Sauerbrei B A, Guo J Z, Cohen J D, Mischiati M, Guo W, Kabra M, Verma N, Mensh B, Branson K and Hantman A W 2020 Cortical pattern generation during dexterous movement is input-driven Nature 577 386–91

[52] Musall S, Kaufman M T, Juavinett A L, Gluf S and Churchland A K 2019 Single-trial neural dynamics are dominated by richly varied movements Nat. Neurosci. 22 167786

[53] Luo T Z, Bondy A G, Gupta D, Elliott V A, Kopec C D and Brody C D 2020 An approach for long-term, multi-probe Neuropixels recordings in unrestrained rats bioRxiv 039305

[54] Juavinett A L, Bekheet G and Churchland A K 2019 Chronically implanted Neuropixels probes enable high-yield recordings in freely moving mice eLife 8 e47188

[55] Paxinos G and Watson C 2007 The rat brain in stereotaxic coordinates (Academic Press/Elsevier)

[56] DiCarlo J J, Lane J W, Hsiao S S and Johnson K O 1996 Marking microelectrode penetrations with fluorescent dyes J. Neurosci. Methods 64 75–81

[57] Fiath R, Kerekes B P, Wittner L, Toth K, Beregszaszi P, Horvath D and Ulbert I 2016 Laminar analysis of the slow wave activity in the somatosensory cortex of anesthetized rats Eur. J. Neurosci. 44 1935–51

[58] Pachitariu M, Steinmetz N, Kadir S, Carandini M and Harris K D 2016 Kilosort: realtime spike-sorting for extracellular electrophysiology with hundreds of channels bioRxiv 061481

[59] Schmitzer-Torbert N, Jackson J, Henze D, Harris K and Redish A D 2005 Quantitative measures of cluster quality for use in extracellular recordings Neuroscience 131 1–11

[60] Gold C, Henze D A, Koch C and Buzsaki G 2006 On the origin of the extracellular action potential waveform: A modeling study J. Neurophysiol. 95 3113–28

[61] Fiath R, Hofer K T, Csikos V, Horvath D, Nanasi T, Toth K, Pothof F, Bohler C, Asplund M, Ruther P and Ulbert I 2018 Long-term recording performance and biocompatibility of chronically implanted cylindrically-shaped, polymer-based neural interfaces Biomed. Tech. (Berl.) 63 301–15

[62] Harris K D and Shepherd G M 2015 The neocortical circuit: themes and variations Nat. Neurosci. 18 170–81

[63] Moffitt M A and McIntyre C C 2005 Model-based analysis of cortical recording with silicon microelectrodes Clin. Neurophysiol. 116 2240–50

[64] Du J, Riedel-Kruse I H, Nawroth J C, Roukes M L, Laurent G and Masmanidis S C 2009 High-resolution three-dimensional extracellular recording of neuronal activity with microfabricated electrode arrays J. Neurophysiol. 101 1671–8

[65] Seymour J P and Kipke D R 2007 Neural probe design for reduced tissue encapsulation in CNS Biomaterials 28 3594–607

[66] Polikov V S, Tresco P A and Reichert W M 2005 Response of brain tissue to chronically implanted neural electrodes J. Neurosci. Methods 148 1–18

[67] Weltman A, Yoo J and Meng E 2016 Flexible, Penetrating Brain Probes Enabled by Advances in Polymer Microfabrication Micromachines 7 180

