## Supplemental Figures and Tables for "Recording site placement on planar silicon-based probes affects neural signal quality: edge sites enhance acute recording performance"

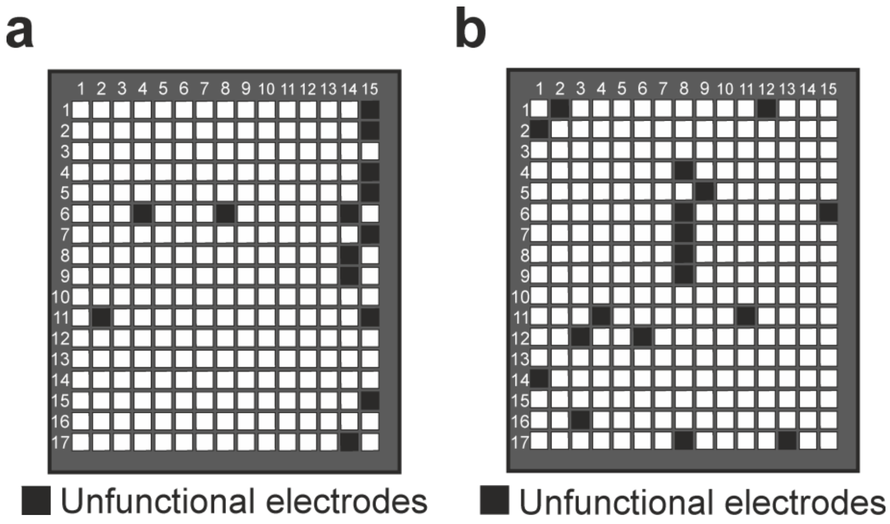


**Supplementary Figure 1.** The position of unfunctional recording sites/channels (black squares) on the 255-channel silicon probe used in the study (a) and on the 255-channel probe which was used to collect the online available data ((b); www.kampff-lab.org/ultra-dense-survey).

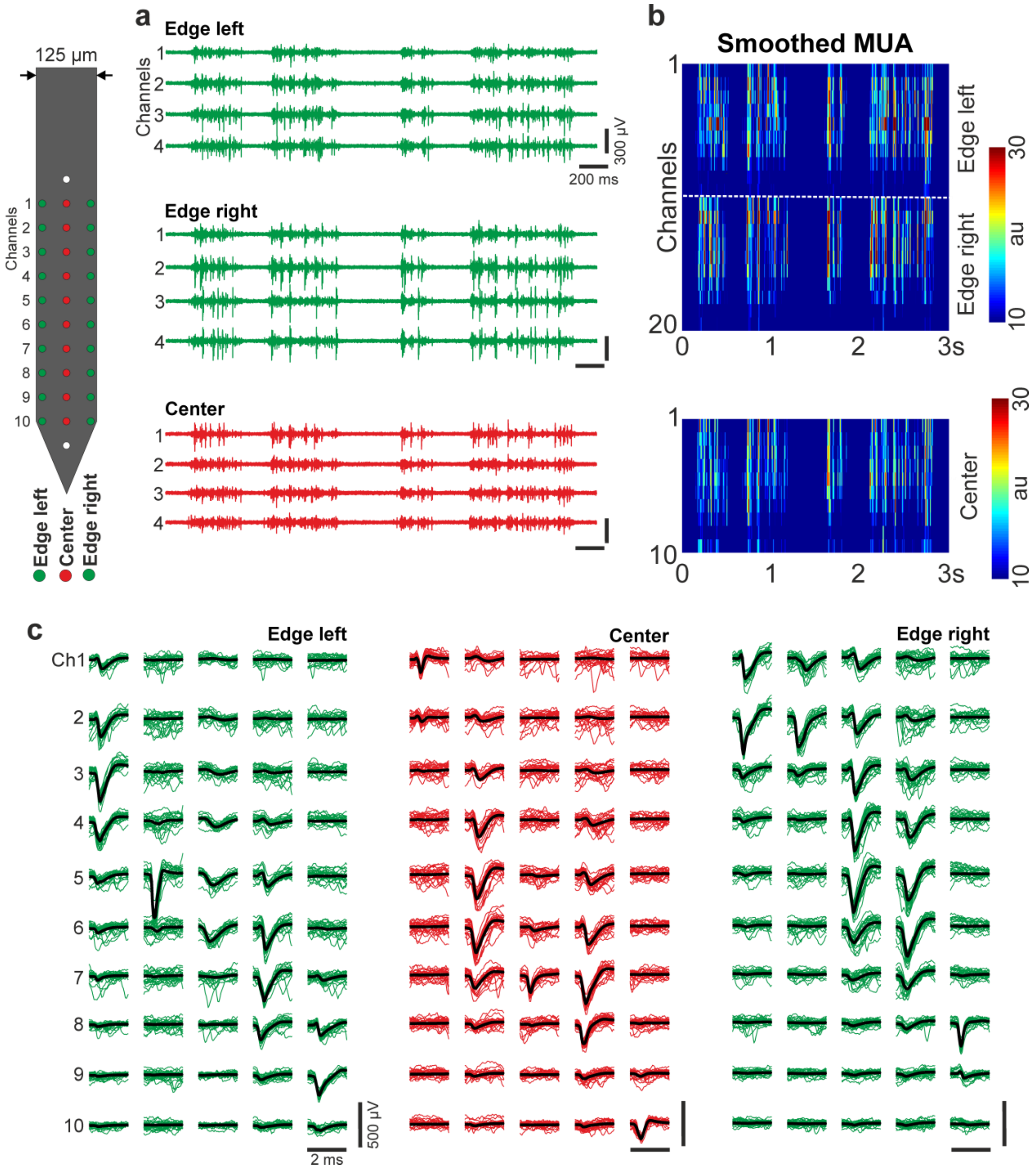


**Supplementary Figure 2.** Representative cortical data obtained with the 32-channel NeuroNexus probe from an anesthetized rat. (a) Examples of three-second-long multiunit activity (MUA; 500-5000 Hz) traces. Four channels for each site position are shown. (b) Rectified and smoothed MUA (50 Hz lowpass filter) recorded on all edge and center channels. The dashed white line separates channels located on the left and right edge of the probe (10 channels/site position; au, arbitrary unit). (c) Five exemplary single units for each site position. For each single unit, twenty-five superposed individual wideband spikes (thin colored lines) and the average spike waveform (thick black line) are shown on ten channels.

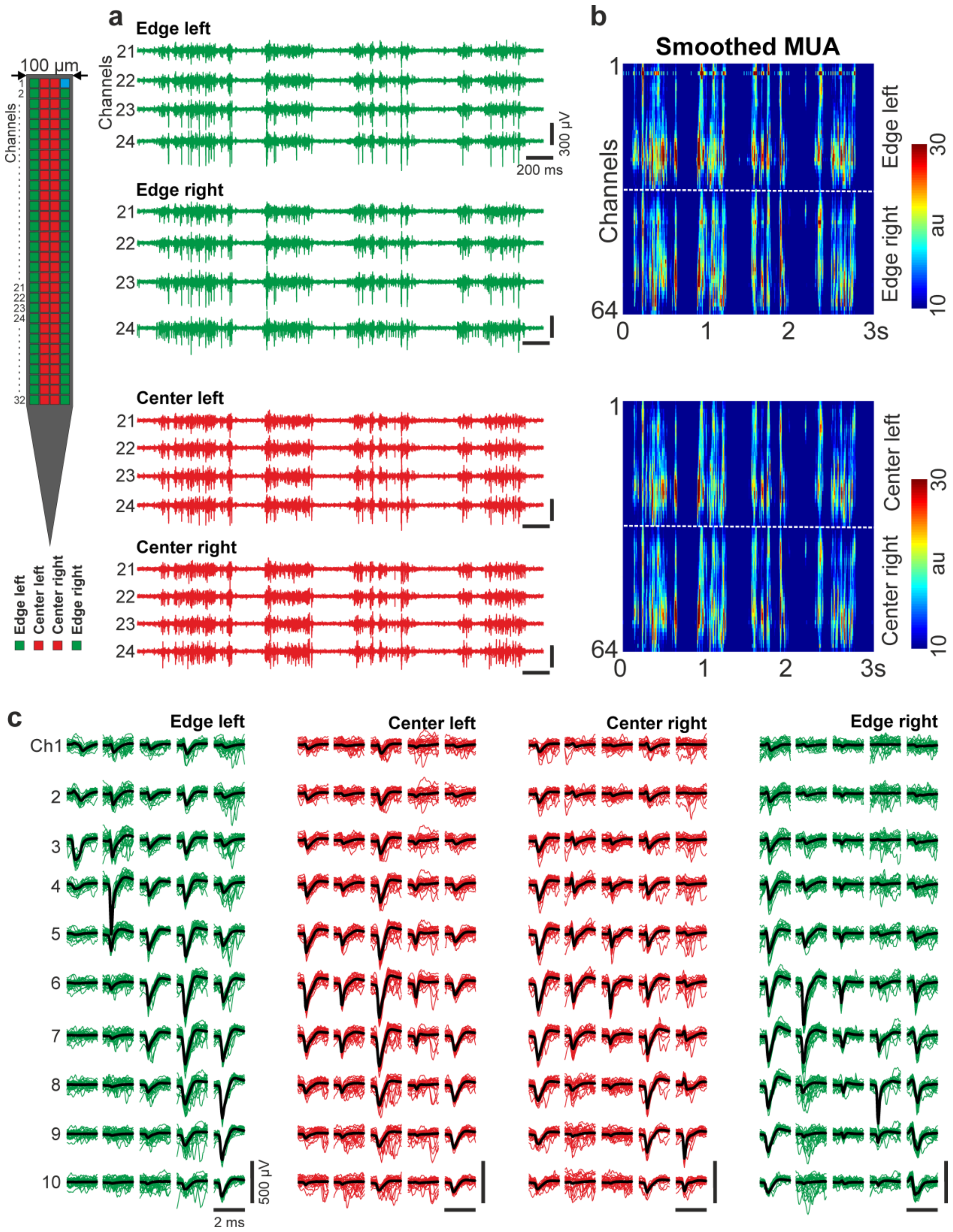


**Supplementary Figure 3.** Representative cortical data obtained with the 128-channel NeuroSeeker probe from an anesthetized rat. (a) Examples of three-second-long multiunit activity (MUA; 500-5000 Hz) traces. Four channels for each site position are shown. (b) Rectified and smoothed MUA (50 Hz lowpass filter) recorded on all edge and center channels. The dashed white lines separate channels located on the left and right side of the probe (32 channels/site position; au, arbitrary unit). (c) Five exemplary single units for each site position. For each single unit, twenty-five superposed individual wideband spikes (thin colored lines) and the average spike waveform (thick black line) are shown on ten adjacent channels.

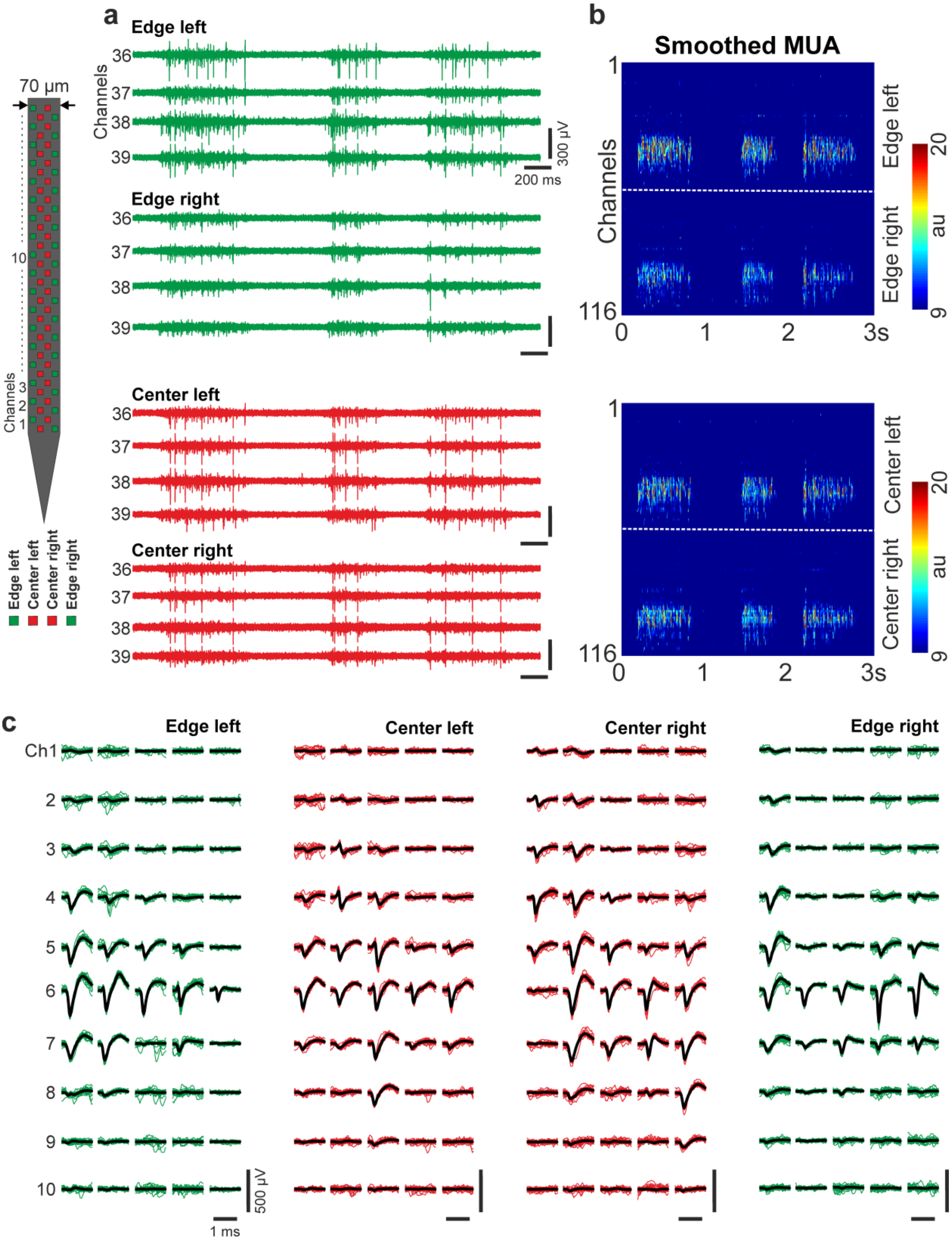


**Supplementary Figure 4.** Representative cortical data obtained with the 70-µm-wide Neuropixels probe from an anesthetized rat. (a) Examples of three-second-long multiunit activity (MUA; 500-5000 Hz) traces. Four channels for each site position are shown. (b) Rectified and smoothed MUA (50 Hz lowpass filter) recorded on all edge and center channels. The dashed white lines separate channels located on the left and right side of the probe (58 channels/site position; au, arbitrary unit). (c) Five exemplary single units for each site position.

For each single unit, twenty-five superposed individual spikes (thin colored lines, AP band) and the average spike waveform (thick black line) are shown on ten adjacent channels.

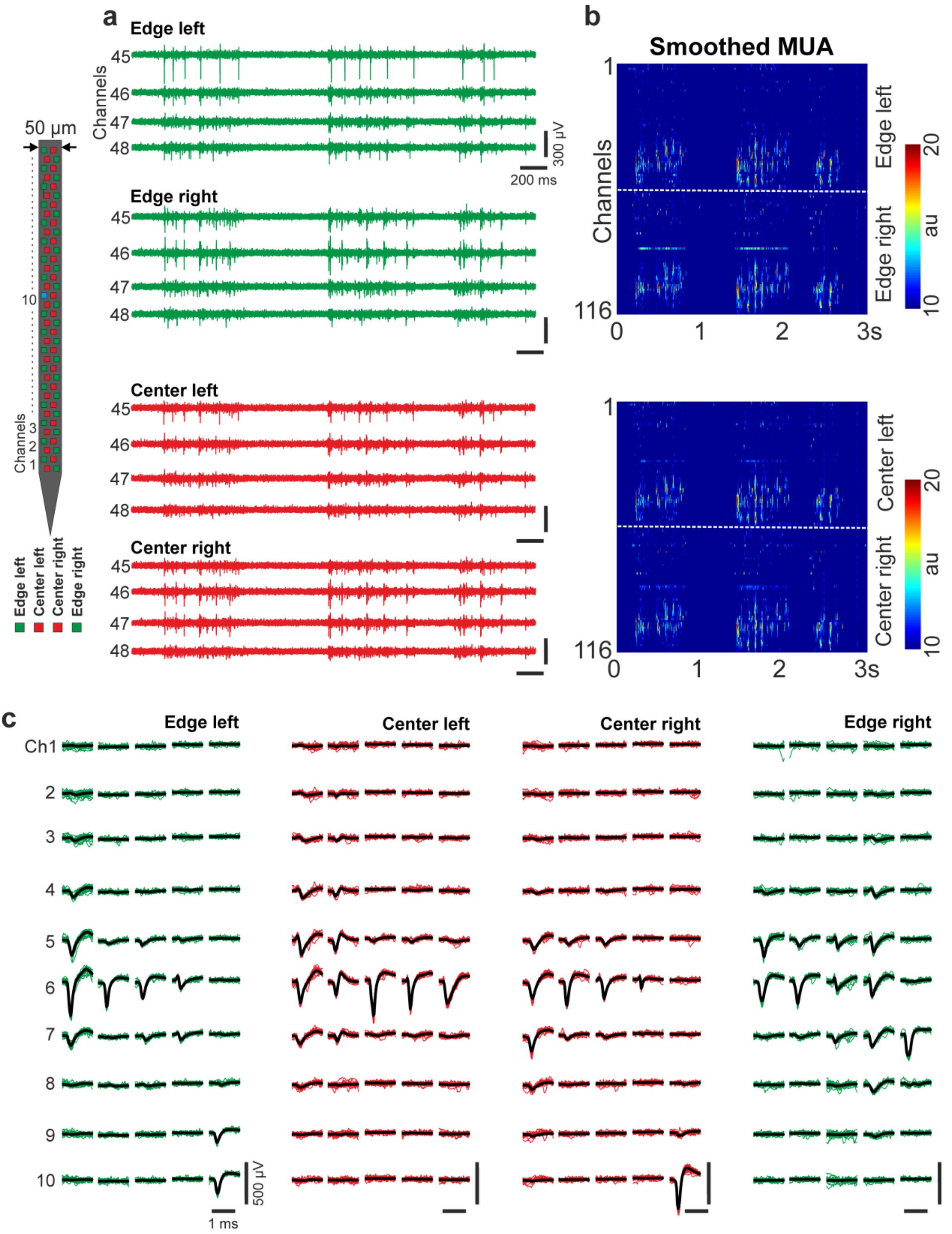


**Supplementary Figure 5.** Representative cortical data obtained with the 50-µm-wide Neuropixels probe from an anesthetized rat. (a) Examples of three-second-long multiunit activity (MUA; 500-5000 Hz) traces. Four channels for each site position are shown. (b) Rectified and smoothed MUA (50 Hz lowpass filter) recorded on all edge and center channels. The dashed white lines separate channels located on the left and right side of the probe (58 channels/site position; au, arbitrary unit). (c) Five exemplary single units for each site position. For each single unit, twenty-five superposed individual spikes (thin colored lines, AP band) and the average spike waveform (thick black line) are shown on ten adjacent channels.

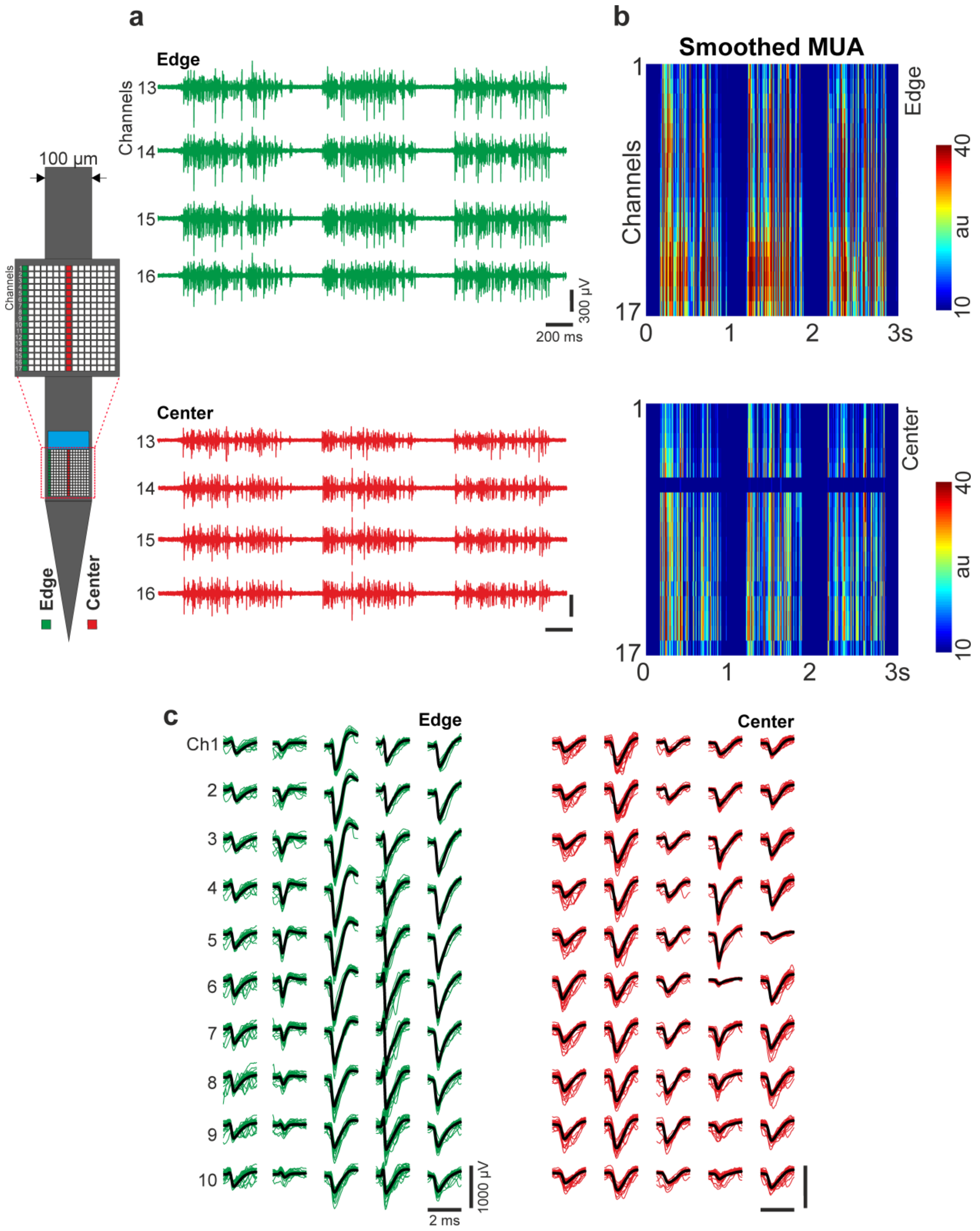


**Supplementary Figure 6.** Representative cortical data obtained with the 255-channel NeuroSeeker probe from an anesthetized rat. (a) Examples of three-second-long multiunit activity (MUA; 500-5000 Hz) traces. Four channels for each site position are shown. (b) Rectified and smoothed MUA (50 Hz lowpass filter) recorded on all left edge and center channels (17 channels/site position; au, arbitrary unit). (c) Five exemplary single units for each site position. For each single unit, twenty-five superposed individual wideband spikes (thin colored lines) and the average spike waveform (thick black line) are shown on ten adjacent channels.

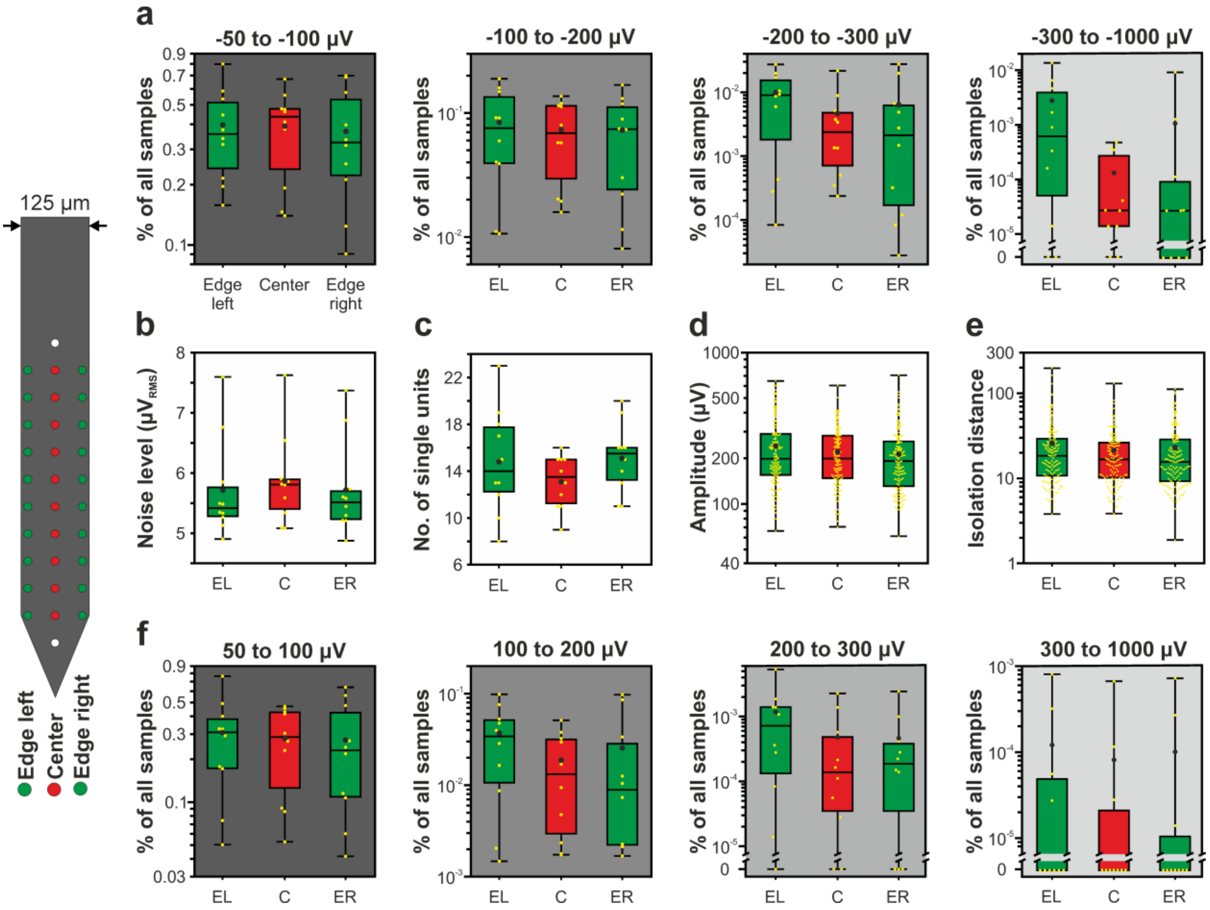


**Supplementary Figure 7.** Boxplots showing the results of the 32-channel NeuroNexus silicon probe for edge (green, both for left and right side) and center (red) sites. (a) Ratio of samples to the total number of samples for each of the four negative amplitude ranges (n = 10 recordings). (b) Estimated *in vivo* noise level. (c) Single unit yield (n = 430). (d) Peak-to-peak amplitude of the averaged single unit spike waveforms. (e) Isolation distance of the single unit clusters. (f) Ratio of samples to the total number of samples for each of the four positive amplitude ranges (n = 10 recordings). All boxplots in the Supplementary material are presented as follows (see also figure 3(c)). The middle line indicates the median, while the boxes correspond to the 25th and 75th percentile. Whiskers mark the minimum and maximum values. The average is depicted with a black dot, while individual values are indicated with smaller yellow dots. Note that most data are plotted on a logarithmic scale.

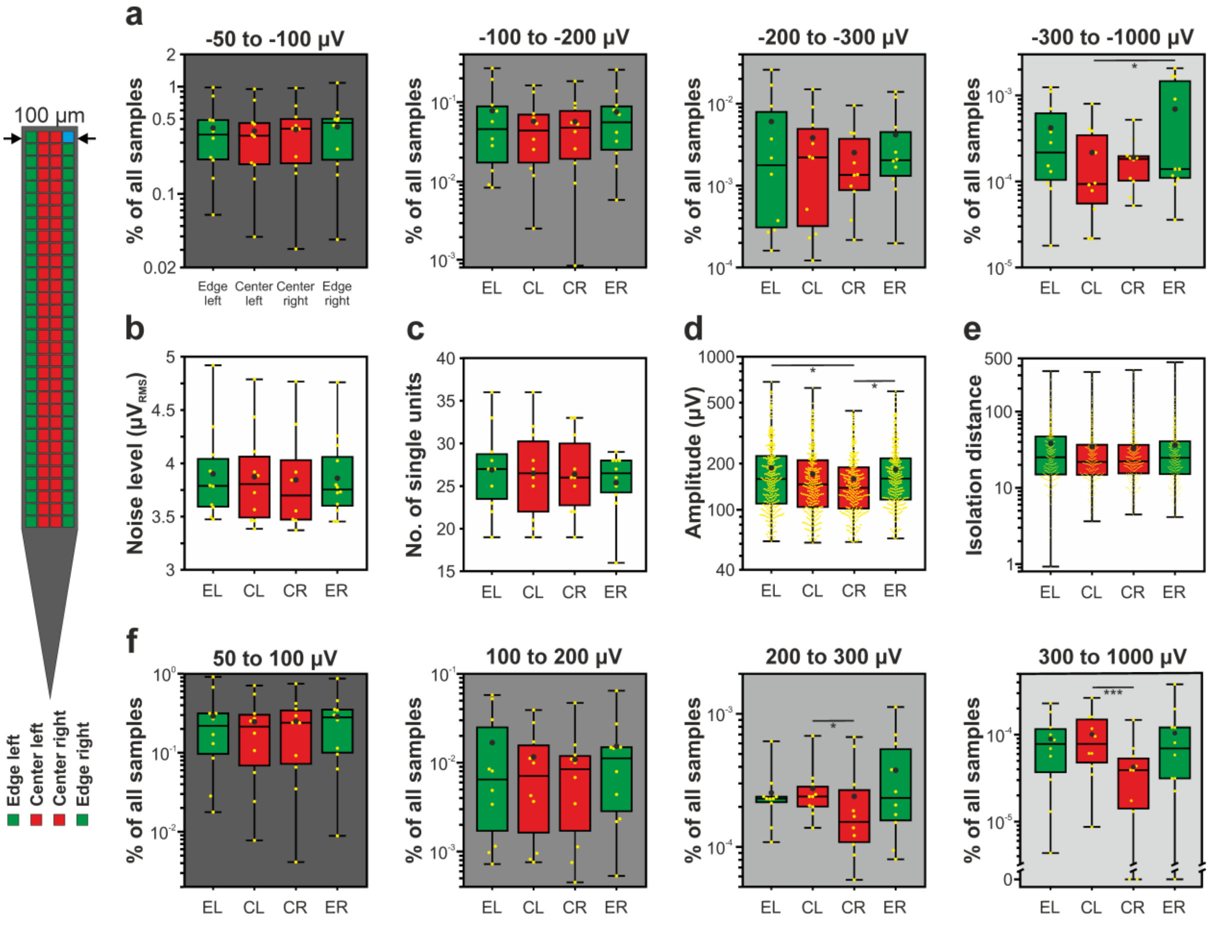


**Supplementary Figure 8.** Boxplots showing the results of the 128-channel NeuroSeeker silicon probe for edge (green) and center (red) sites, separated by left and right side. (a) Ratio of samples to the total number of samples for each of the four negative amplitude ranges (n = 10 recordings). (b) Estimated *in vivo* noise level. (c) Single unit yield (n = 1052). (d) Peak-to-peak amplitude of the averaged single unit spike waveforms. (e) Isolation distance of the single unit clusters. (f) Ratio of samples to the total number of samples for each of the four positive amplitude ranges (n = 10 recordings). Note that most data are plotted on a logarithmic scale. * $p < 0.05$; *** $p < 0.001$.

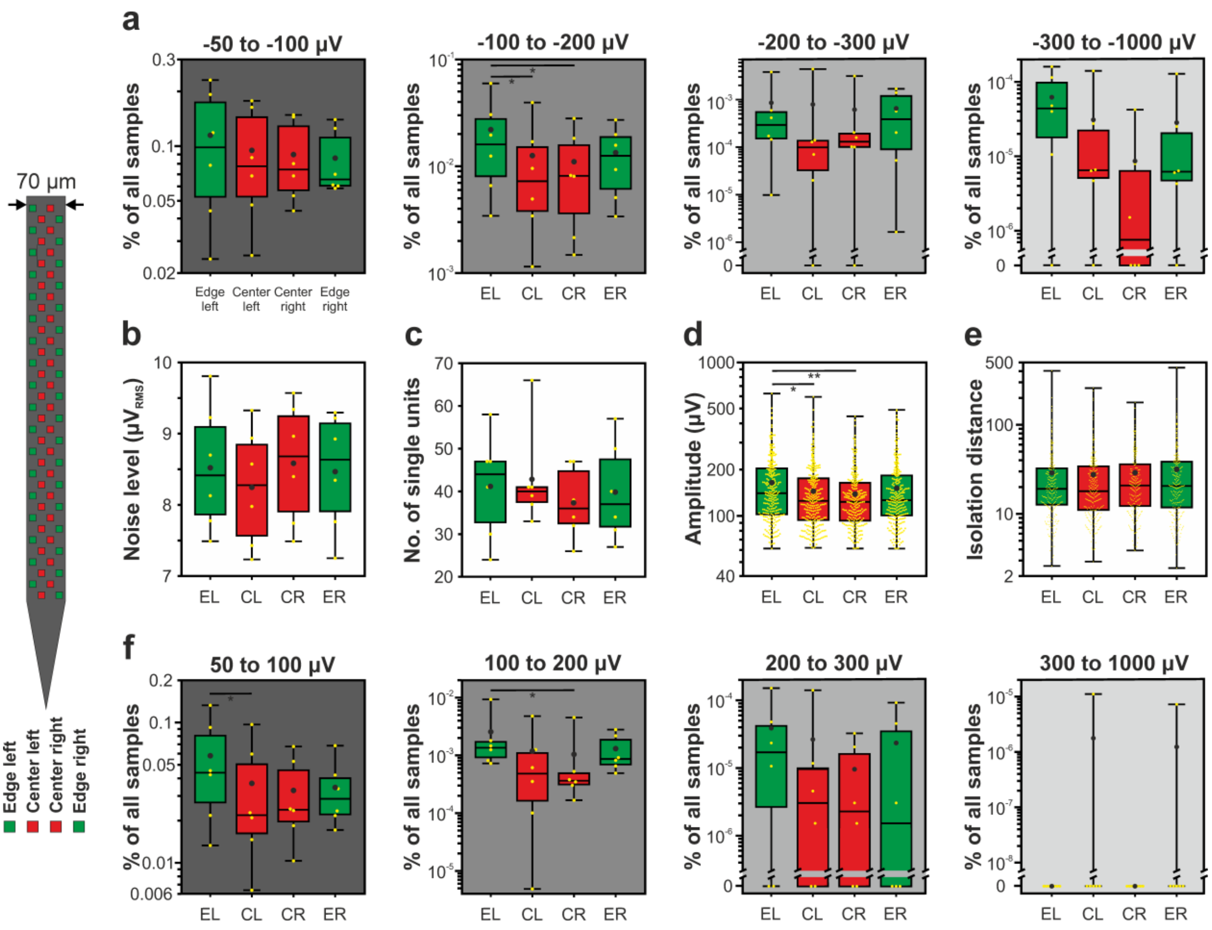


**Supplementary Figure 9.** Boxplots showing the results of the 70-µm-wide Neuropixels probe for edge (green) and center (red) sites, separated by left and right side. (a) Ratio of samples to the total number of samples for each of the four negative amplitude ranges (n = 6 recordings). (b) Estimated *in vivo* noise level. (c) Single unit yield (n = 967). (d) Peak-to-peak amplitude of the averaged single unit spike waveforms. (e) Isolation distance of the single unit clusters. (f) Ratio of samples to the total number of samples for each of the four positive amplitude ranges (n = 6 recordings). Note that most data are plotted on a logarithmic scale. * $p < 0.05$; ** $p < 0.01$.

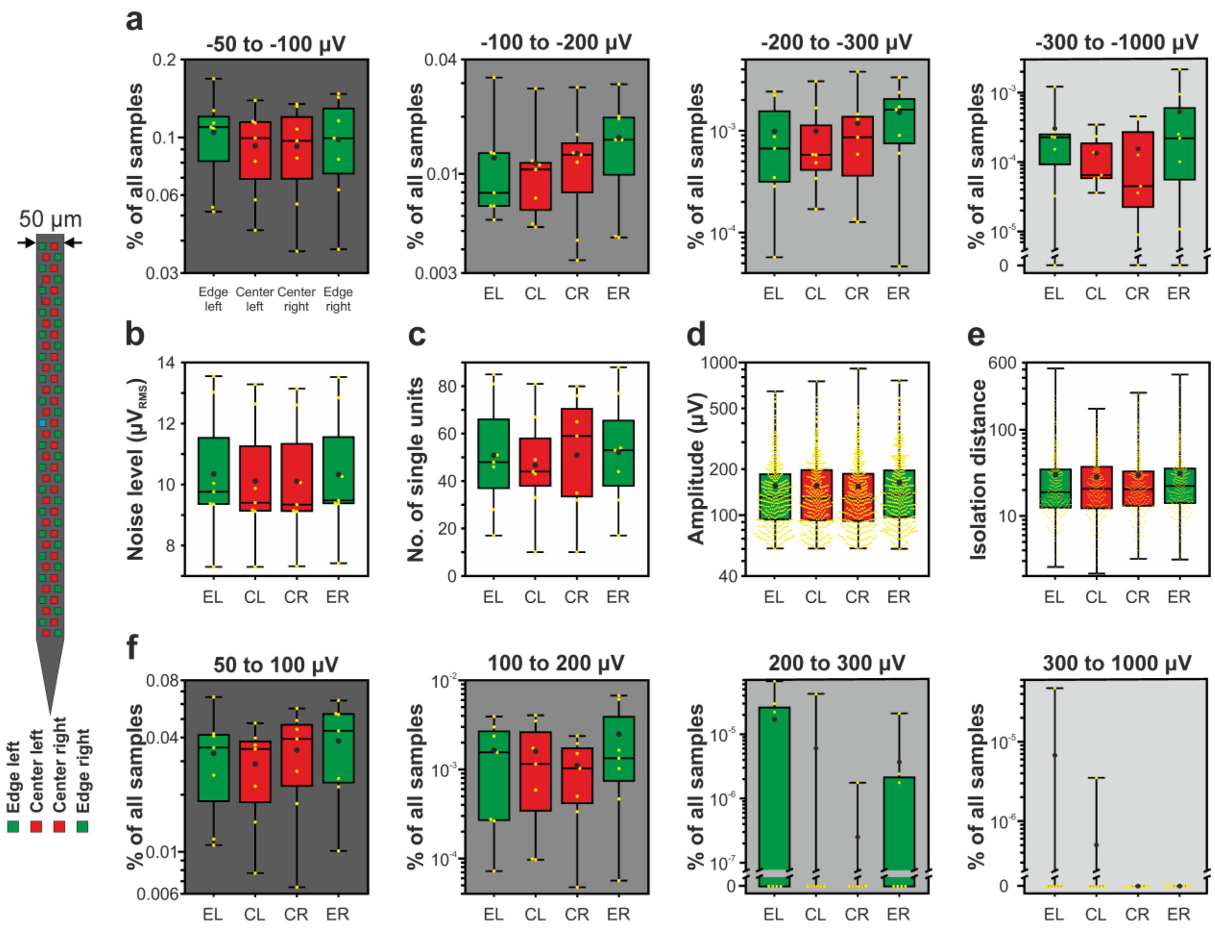


**Supplementary Figure 10.** Boxplots showing the results of the 50-µm-wide Neuropixels probe for edge (green) and center (red) sites, separated by left and right side. (a) Ratio of samples to the total number of samples for each of the four negative amplitude ranges (n = 7 recordings). (b) Estimated *in vivo* noise level. (c) Single unit yield (n = 1405). (d) Peak-to-peak amplitude of the averaged single unit spike waveforms. (e) Isolation distance of the single unit clusters. (f) Ratio of samples to the total number of samples for each of the four positive amplitude ranges (n = 7 recordings). Note that most data are plotted on a logarithmic scale.

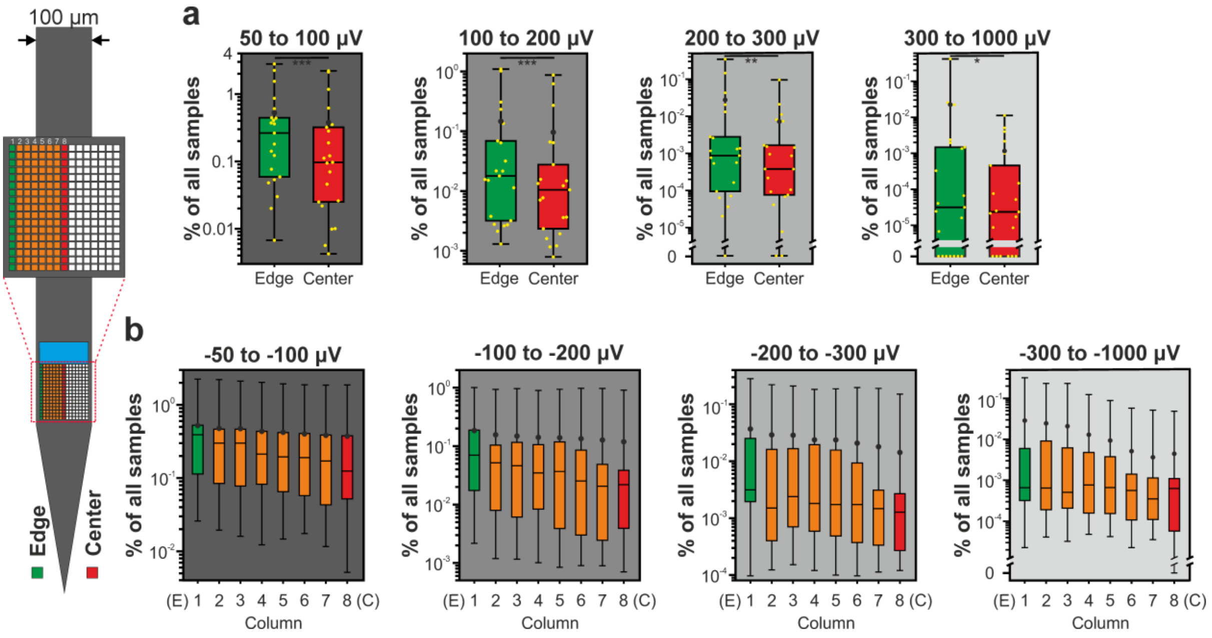


**Supplementary Figure 11.** Boxplots showing the results of the 255-channel NeuroSeeker probe for edge (green) and center (red). (a) Ratio of samples to the total number of samples for each of the four positive amplitude ranges (n = 21 recordings). (b) Ratio of samples to the total number of samples for each of the four negative amplitude ranges (n = 21 recordings) shown for edge sites (green), center sites (red) and columns of sites located between these (orange; columns 2-7; see inset on the left side of the figure). Note that data are plotted on a logarithmic scale. * $p < 0.05$; ** $p < 0.01$; *** $p < 0.001$.

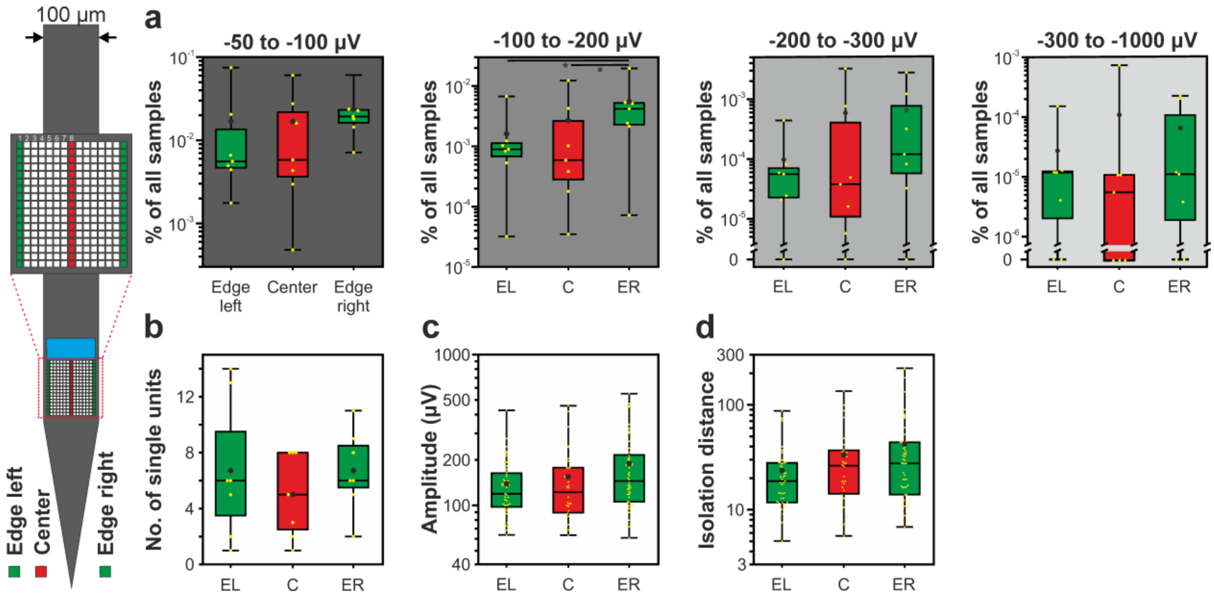


**Supplementary Figure 12.** Boxplots showing the results of the online available 255-channel silicon probe data (www.kampff-lab.org/ultra-dense-survey) for edge (green) and center (red) sites. (a) Ratio of samples to the total number of samples for each of the four amplitude ranges (n = 7 recordings). (b) Single unit yield (n = 129). (c) Peak-to-peak amplitude of the averaged single unit spike waveforms. (d) Isolation distance of the single unit clusters. Note that most data are plotted on a logarithmic scale. * $p < 0.05$.

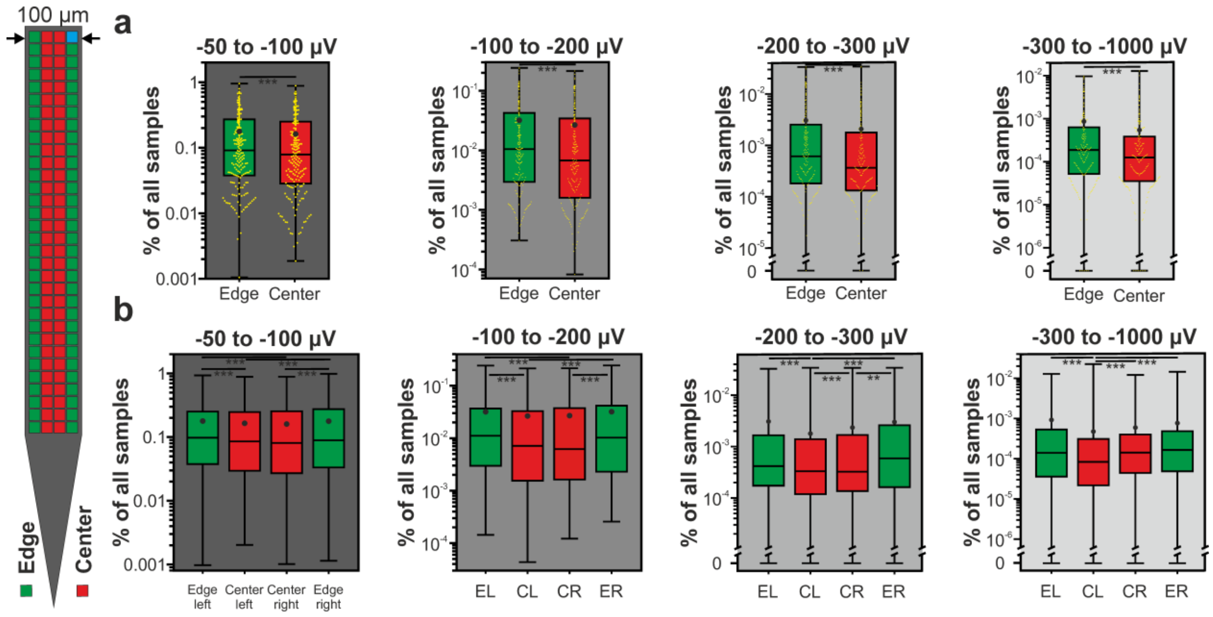


**Supplementary Figure 13.** Boxplots showing the results of the larger dataset (n = 186 recordings) obtained with the 128-channel NeuroSeeker silicon probe for edge (green) and center (red) sites. Ratio of samples to the total number of samples for each of the four negative amplitude ranges for pooled data (a) and data separated by left and right side (b). Note that data are plotted on a logarithmic scale. ** $p < 0.01$; *** $p < 0.001$.

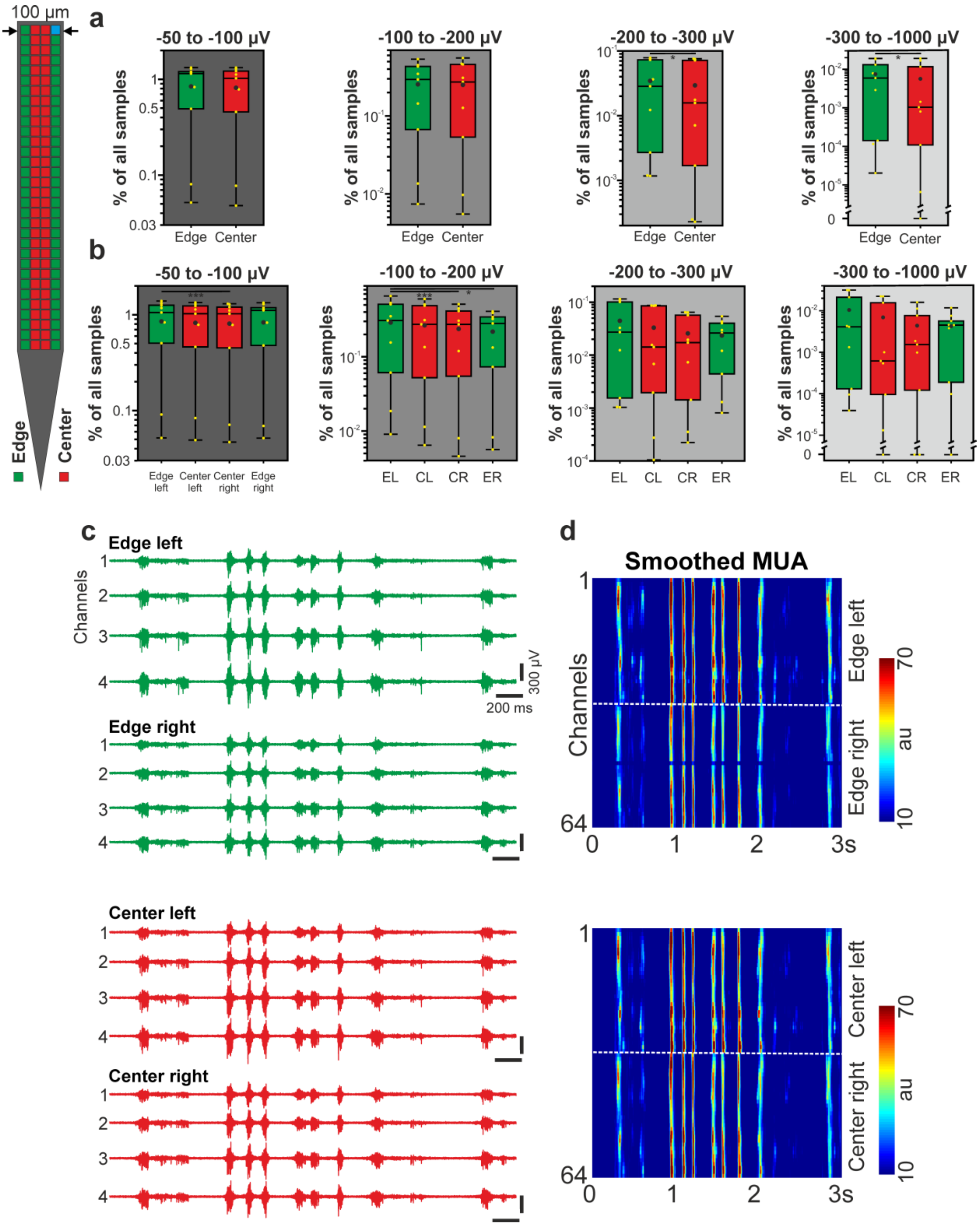


**Supplementary Figure 14.** Boxplots showing the results of the thalamic recordings (n = 9) obtained with the 128-channel NeuroSeeker silicon probe for edge (green) and center (red) sites. Ratio of samples to the total number of samples for each of the four negative amplitude ranges for pooled data (a) and data separated by left and right side (b). Note that data are plotted on a logarithmic scale. (c) Examples of three-second-long thalamic multiunit activity (MUA; 500-5000 Hz) traces. Four channels for each site position are shown. (d) Rectified and smoothed MUA (50 Hz lowpass filter) recorded on all edge and center channels. The dashed white lines separate channels located on the left and right side of the probe (32 channels/site position; au, arbitrary unit). * $p < 0.05$; *** $p < 0.001$.

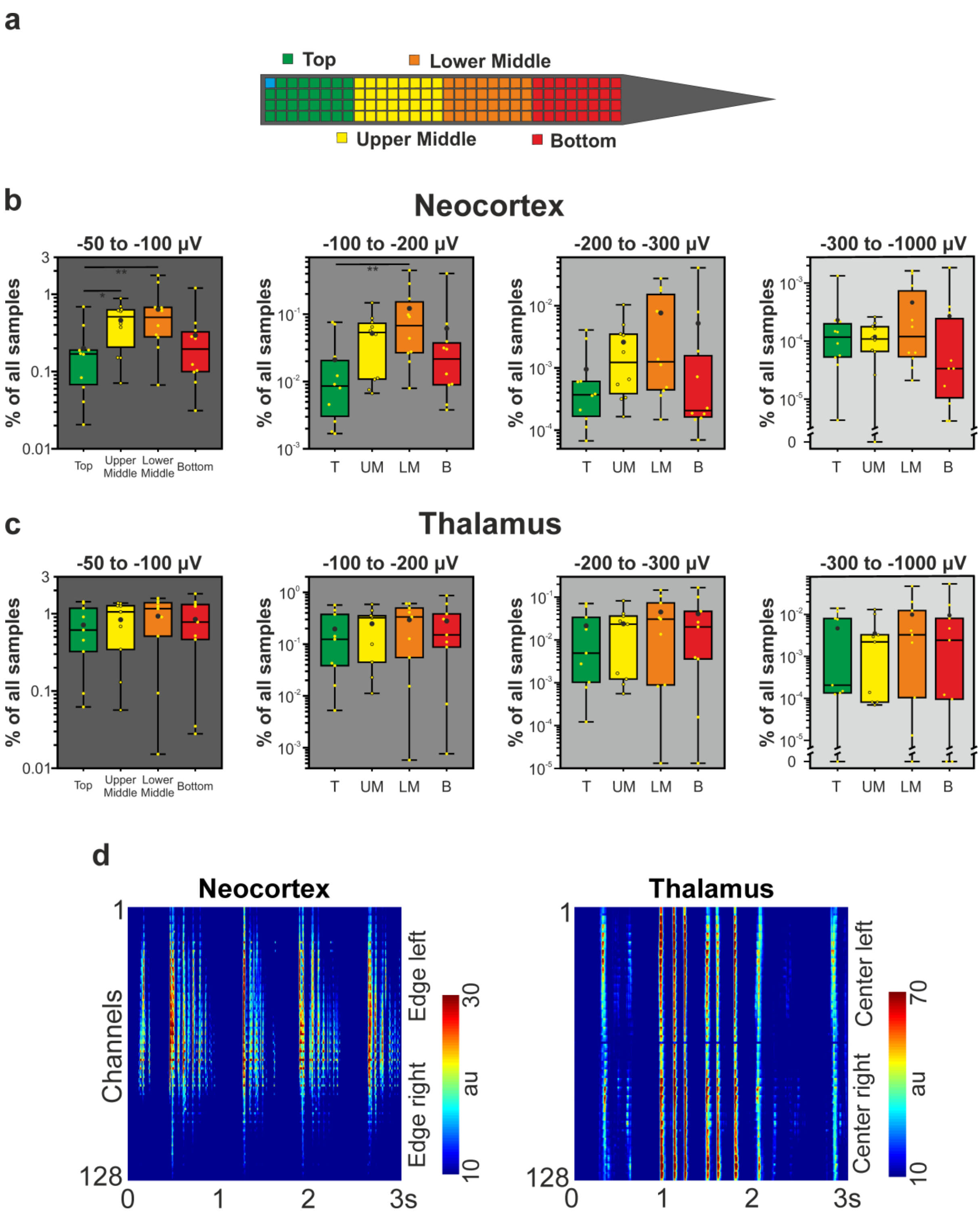


**Supplementary Figure 15.** Results of the analysis based on longitudinal site position. (a) The recording sites of the 128-channel probe were grouped according to their longitudinal position (32-channel/site position). Four site groups were constructed with the bottom sites located closest to the probe tip. (b-c) Ratio of samples to the total number of samples for each of the four negative amplitude ranges for the (b) cortical data (n = 10 recordings) and for the (c) thalamic data (n = 9 recordings), separated by longitudinal site position. Note that data are plotted on a logarithmic scale. (d) Three-second-long rectified and smoothed (50 Hz lowpass filter) multiunit activity recorded in the cortex (left) and in the thalamus (right). Note that, compared to the thalamus, cortical spiking activity is usually not recorded simultaneously with all sites. * $p < 0.05$; ** $p < 0.01$.

| Probe type | No. of recording sites | No. of sites in separated recordings | No. of analysed recordings | No. of rats | No. of penetrations | Average recording length (min) | Reference |
|---|---|---|---|---|---|---|---|
| 255-channel NeuroSeeker probe | 255 | 17 | 7 | 3 | 3 | 27 | Dimitriadis et al., 2018, bioRxiv 275818 |

**Supplementary Table 1.** Details of the experiments and recordings of the 255-channel silicon probe data available online (www.kampff-lab.org/ultra-dense-survey). Cortical recordings with the following identifiers were used in the analysis: Co1, Co2, Co3, Co5, CoP1, CoP2, CoP3.

| Probe type | No. of recording sites | No. of sites in separated recordings | No. of analysed recordings | No. of rats | No. of penetrations | Type of anesthesia | Average recording length (min) |
|---|---|---|---|---|---|---|---|
| 128-channel NeuroSeeker probe | 128 | 32 | 9 | 3 | 3 | Ketamine/xylazine | 12 |

**Supplementary Table 2.** Details of the thalamic experiments and recordings of the 128-channel probe.

| Probe type | No. of excluded single units | Total number of single units | % of units excluded |
|---|---|---|---|
| 32-channel NeuroNexus probe | 6 | 430 | 1.38 |
| 128-channel NeuroSeeker probe | 156 | 1052 | 12.91 |
| Neuropixels probe (70 µm thickness) | 61 | 967 | 5.93 |
| Neuropixels probe (50 µm thickness) | 95 | 1405 | 6.33 |
| 255-channel NeuroSeeker probe | 27 | 599 | 4.31 |

**Supplementary Table 3.** Number and ratio of single units excluded from the analysis due to poor cluster quality.

| | 32-channel NN probe | | |
|---|---|---|---|
| | Edge Left | Center | Edge Right |
| −50 to −100 μV (% of all samples) | **0.398 ± 0.200** | 0.391 ± 0.179 | 0.369 ± 0.219 |
| −100 to −200 μV (% of all samples) | **$8.40 \times 10^{-2} \pm 6.34 \times 10^{-2}$** | $7.34 \times 10^{-2} \pm 4.59 \times 10^{-2}$ | $7.27 \times 10^{-2} \pm 5.49 \times 10^{-2}$ |
| −200 to −300 μV (% of all samples) | **$9.93 \times 10^{-3} \pm 9.12 \times 10^{-3}$** | $4.66 \times 10^{-3} \pm 6.55 \times 10^{-3}$ | $6.38 \times 10^{-3} \pm 9.58 \times 10^{-3}$ |
| −300 to −1000 μV (% of all samples) | **$2.79 \times 10^{-3} \pm 4.42 \times 10^{-3}$** | $1.32 \times 10^{-4} \pm 1.87 \times 10^{-4}$ | $1.06 \times 10^{-3} \pm 2.86 \times 10^{-3}$ |
| 50 to 100 μV (% of all samples) | **0.307 ± 0.213** | 0.280 ± 0.162 | 0.273 ± 0.214 |
| 100 to 200 μV (% of all samples) | **$3.70 \times 10^{-2} \pm 3.22 \times 10^{-2}$** | $1.87 \times 10^{-2} \pm 1.77 \times 10^{-2}$ | $2.55 \times 10^{-2} \pm 3.61 \times 10^{-2}$ |
| 200 to 300 μV (% of all samples) | **$1.18 \times 10^{-3} \pm 1.60 \times 10^{-3}$** | $4.79 \times 10^{-4} \pm 7.58 \times 10^{-4}$ | $4.63 \times 10^{-4} \pm 7.53 \times 10^{-3}$ |
| 300 to 1000 μV (% of all samples) | **$1.21 \times 10^{-4} \pm 2.60 \times 10^{-4}$** | $8.10 \times 10^{-5} \pm 2.09 \times 10^{-4}$ | $1.00 \times 10^{-4} \pm 2.34 \times 10^{-4}$ |
| No. of single units | 14.80 ± 3.73 | 13.10 ± 2.28 | **15.10 ± 2.99** |
| Amplitude of single units (μV) | **240.46 ± 127.62** | 220.11 ± 97.39 | 211.90 ± 105.30 |
| Isolation distance | **25.84 ± 26.90** | 21.35 ± 17.3 | 22.94 ± 20.43 |
| Noise level ($\mu V_{RMS}$) | 5.72 ± 0.83 | **5.87 ± 0.75** | 5.72 ± 0.79 |

**Supplementary Table 4.** Mean ± standard deviation of the calculated features for the 32-channel NeuroNexus probe. For each feature, the largest mean value is indicated in bold.

| | 128-channel NS probe | | | |
|---|---|---|---|---|
| | Edge Left | Center Left | Center Right | Edge Right |
| −50 to −100 μV (% of all samples) | 0.412 ± 0.296 | 0.387 ± 0.283 | 0.399 ± 0.276 | **0.422 ± 0.295** |
| −100 to −200 μV (% of all samples) | **$7.81 \times 10^{-2} \pm 8.68 \times 10^{-2}$** | $5.74 \times 10^{-2} \pm 5.34 \times 10^{-2}$ | $5.69 \times 10^{-2} \pm 5.44 \times 10^{-2}$ | $7.53 \times 10^{-2} \pm 7.53 \times 10^{-2}$ |
| −200 to −300 μV (% of all samples) | **$6.04 \times 10^{-3} \pm 8.81 \times 10^{-3}$** | $3.83 \times 10^{-3} \pm 4.84 \times 10^{-3}$ | $2.53 \times 10^{-3} \pm 2.87 \times 10^{-3}$ | $4.18 \times 10^{-3} \pm 4.85 \times 10^{-3}$ |
| −300 to −1000 μV (% of all samples) | $4.19 \times 10^{-4} \pm 4.46 \times 10^{-4}$ | $2.17 \times 10^{-4} \pm 2.49 \times 10^{-4}$ | $1.91 \times 10^{-4} \pm 1.39 \times 10^{-4}$ | **$6.93 \times 10^{-4} \pm 8.10 \times 10^{-4}$** |
| 50 to 100 μV (% of all samples) | **0.292 ± 0.291** | 0.247 ± 0.231 | 0.250 ± 0.225 | 0.287 ± 0.215 |
| 100 to 200 μV (% of all samples) | **$1.69 \times 10^{-2} \pm 2.20 \times 10^{-2}$** | $1.16 \times 10^{-2} \pm 1.31 \times 10^{-2}$ | $1.09 \times 10^{-2} \pm 1.38 \times 10^{-2}$ | $1.54 \times 10^{-2} \pm 1.92 \times 10^{-2}$ |
| 200 to 300 μV (% of all samples) | $2.55 \times 10^{-4} \pm 1.40 \times 10^{-4}$ | $2.75 \times 10^{-4} \pm 1.53 \times 10^{-4}$ | $2.40 \times 10^{-4} \pm 2.12 \times 10^{-4}$ | **$3.77 \times 10^{-4} \pm 3.35 \times 10^{-4}$** |
| 300 to 1000 μV (% of all samples) | $8.93 \times 10^{-5} \pm 7.25 \times 10^{-5}$ | $1.01 \times 10^{-4} \pm 7.73 \times 10^{-5}$ | $4.26 \times 10^{-5} \pm 4.37 \times 10^{-5}$ | **$1.04 \times 10^{-4} \pm 1.13 \times 10^{-4}$** |
| No. of single units | **26.90 ± 5.07** | 26.50 ± 5.52 | 26.40 ± 4.77 | 25.40 ± 3.84 |
| Amplitude of single units (μV) | **186.90 ± 108.42** | 170.63 ± 92.56 | 158.65 ± 77.20 | 182.85 ± 95.12 |
| Isolation distance | **38.06 ± 40.68** | 34.41 ± 39.16 | 32.63 ± 38.00 | 36.04 ± 43.68 |
| Noise level ($\mu V_{RMS}$) | **3.90 ± 0.45** | 3.87 ± 0.46 | 3.85 ± 0.44 | 3.86 ± 0.42 |

**Supplementary Table 5.** Mean ± standard deviation of the calculated features for the 128-channel NeuroSeeker probe.

| | 70 µm NP probe | | | |
|---|---|---|---|---|
| | Edge Left | Center Left | Center Right | Edge Right |
| −50 to −100 µV (% of all samples) | **0.114 ± 0.082** | 0.085 ± 0.036 | 0.089 ± 0.045 | 0.094 ± 0.062 |
| −100 to −200 µV (% of all samples) | **$2.17 \times 10^{-2} \pm 2.05 \times 10^{-2}$** | $1.32 \times 10^{-2} \pm 9.04 \times 10^{-3}$ | $1.09 \times 10^{-2} \pm 1.01 \times 10^{-2}$ | $1.24 \times 10^{-2} \pm 1.41 \times 10^{-2}$ |
| −200 to −300 µV (% of all samples) | **$8.54 \times 10^{-4} \pm 1.46 \times 10^{-3}$** | $6.50 \times 10^{-4} \pm 7.22 \times 10^{-4}$ | $6.14 \times 10^{-4} \pm 1.23 \times 10^{-3}$ | $7.90 \times 10^{-4} \pm 1.76 \times 10^{-3}$ |
| −300 to −1000 µV (% of all samples) | **$6.20 \times 10^{-5} \pm 6.24 \times 10^{-5}$** | $2.83 \times 10^{-5} \pm 4.96 \times 10^{-5}$ | $8.60 \times 10^{-6} \pm 1.67 \times 10^{-5}$ | $3.09 \times 10^{-5} \pm 5.44 \times 10^{-5}$ |
| 50 to 100 µV (% of all samples) | **$5.80 \times 10^{-2} \pm 4.58 \times 10^{-2}$** | $3.69 \times 10^{-2} \pm 3.46 \times 10^{-2}$ | $3.28 \times 10^{-2} \pm 2.22 \times 10^{-2}$ | $3.45 \times 10^{-2} \pm 1.90 \times 10^{-2}$ |
| 100 to 200 µV (% of all samples) | **$2.57 \times 10^{-3} \pm 3.34 \times 10^{-3}$** | $1.19 \times 10^{-3} \pm 1.83 \times 10^{-3}$ | $1.05 \times 10^{-3} \pm 1.72 \times 10^{-3}$ | $1.31 \times 10^{-3} \pm 9.48 \times 10^{-4}$ |
| 200 to 300 µV (% of all samples) | **$3.99 \times 10^{-5} \pm 5.94 \times 10^{-5}$** | $2.71 \times 10^{-5} \pm 5.77 \times 10^{-5}$ | $9.85 \times 10^{-6} \pm 1.40 \times 10^{-5}$ | $2.41 \times 10^{-5} \pm 3.92 \times 10^{-5}$ |
| 300 to 1000 µV (% of all samples) | 0 | **$1.85 \times 10^{-6} \pm 4.54 \times 10^{-6}$** | 0 | $1.25 \times 10^{-6} \pm 3.07 \times 10^{-6}$ |
| No. of single units | 41.17 ± 12.42 | **42.83 ± 11.74** | 37.33 ± 8.43 | 39.83 ± 11.62 |
| Amplitude of single units (µV) | **163.57 ± 85.77** | 143.77 ± 74.13 | 137.68 ± 65.27 | 150.58 ± 79.36 |
| Isolation distance | 28.92 ± 36.04 | 27.60 ± 32.32 | 28.98 ± 26.79 | **31.63 ± 38.03** |
| Noise level ($\mu V_{RMS}$) | 8.52 ± 0.89 | 8.24 ± 0.84 | **8.58 ± 0.85** | 8.46 ± 0.83 |

**Supplementary Table 6.** Mean ± standard deviation of the calculated features for the 70-µm-wide Neuropixels probe.

| | 50 μm NP probe | | | |
|---|---|---|---|---|
| | Edge Left | Center Left | Center Right | Edge Right |
| −50 to −100 μV (% of all samples) | **0.104 ± 0.041** | 0.093 ± 0.034 | 0.092 ± 0.037 | 0.098 ± 0.041 |
| −100 to −200 μV (% of all samples) | $1.21 \times 10^{-2} \pm 9.26 \times 10^{-3}$ | $1.14 \times 10^{-2} \pm 7.82 \times 10^{-3}$ | $1.28 \times 10^{-2} \pm 8.31 \times 10^{-3}$ | $\mathbf{1.55 \times 10^{-2} \pm 8.86 \times 10^{-3}}$ |
| −200 to −300 μV (% of all samples) | $9.76 \times 10^{-4} \pm 9.48 \times 10^{-4}$ | $9.77 \times 10^{-4} \pm 1.03 \times 10^{-3}$ | $1.17 \times 10^{-3} \pm 1.25 \times 10^{-3}$ | $\mathbf{1.50 \times 10^{-3} \pm 1.11 \times 10^{-3}}$ |
| −300 to −1000 μV (% of all samples) | $3.00 \times 10^{-4} \pm 4.10 \times 10^{-4}$ | $1.32 \times 10^{-4} \pm 1.14 \times 10^{-4}$ | $1.53 \times 10^{-4} \pm 1.93 \times 10^{-4}$ | $\mathbf{5.20 \times 10^{-4} \pm 7.78 \times 10^{-4}}$ |
| 50 to 100 μV (% of all samples) | $3.32 \times 10^{-2} \pm 1.92 \times 10^{-2}$ | $2.91 \times 10^{-2} \pm 1.46 \times 10^{-2}$ | $3.45 \times 10^{-2} \pm 1.81 \times 10^{-2}$ | $\mathbf{3.86 \times 10^{-2} \pm 1.97 \times 10^{-2}}$ |
| 100 to 200 μV (% of all samples) | $1.65 \times 10^{-3} \pm 1.53 \times 10^{-3}$ | $1.61 \times 10^{-3} \pm 1.61 \times 10^{-3}$ | $1.11 \times 10^{-3} \pm 8.78 \times 10^{-4}$ | $\mathbf{2.51 \times 10^{-3} \pm 2.78 \times 10^{-4}}$ |
| 200 to 300 μV (% of all samples) | $\mathbf{1.65 \times 10^{-5} \pm 2.54 \times 10^{-5}}$ | $6.09 \times 10^{-6} \pm 1.61 \times 10^{-5}$ | $2.54 \times 10^{-7} \pm 6.71 \times 10^{-7}$ | $3.64 \times 10^{-6} \pm 7.86 \times 10^{-6}$ |
| 300 to 1000 μV (% of all samples) | $\mathbf{6.85 \times 10^{-6} \pm 1.81 \times 10^{-5}}$ | $5.07 \times 10^{-7} \pm 1.34 \times 10^{-6}$ | 0 | 0 |
| No. of single units | 50.86 ± 25.08 | 46.71 ± 22.91 | 51.00 ± 25.9 | **52.14 ± 24.55** |
| Amplitude of single units (μV) | 155.24 ± 91.12 | 155.96 ± 92.13 | 154.06 ± 93.45 | **164.06 ± 100.24** |
| Isolation distance | 30.12 ± 39.29 | 28.30 ± 24.26 | 29.72 ± 32.68 | **31.28 ± 36.19** |
| Noise level ($\mu V_{RMS}$) | **10.34 ± 2.20** | 10.11 ± 2.11 | 10.10 ± 2.07 | 10.33 ± 2.14 |

**Supplementary Table 7.** Mean ± standard deviation of the calculated features for the 50-μm-wide Neuropixels probe.

| | 255-channel NS probe | |
|---|---|---|
| | Edge | Center |
| 50 to 100 μV (% of all samples) | **0.527 ± 0.757** | 0.376 ± 0.653 |
| 100 to 200 μV (% of all samples) | **0.147 ± 0.314** | 0.096 ± 0.228 |
| 200 to 300 μV (% of all samples) | **$2.76 \times 10^{-2} \pm 7.94 \times 10^{-2}$** | $7.19 \times 10^{-3} \pm 2.09 \times 10^{-2}$ |
| 300 to 1000 μV (% of all samples) | **$2.24 \times 10^{-2} \pm 9.03 \times 10^{-2}$** | $1.16 \times 10^{-3} \pm 2.70 \times 10^{-3}$ |

**Supplementary Table 8.** Mean ± standard deviation of the amplitude distribution of the filtered potential in the four positive amplitude range for the 255-channel NeuroSeeker probe.

| | 255-channel NS probe | | | |
|---|---|---|---|---|
| | Column 1 (Edge) | Column 2 | Column 3 | Column 4 |
| −50 to −100 μV (% of all samples) | **0.521 ± 0.631** | 0.477 ± 0.614 | 0.473 ± 0.62 | 0.434 ± 0.596 |
| −100 to −200 μV (% of all samples) | **0.185 ± 0.289** | 0.156 ± 0.273 | 0.149 ± 0.264 | 0.141 ± 0.259 |
| −200 to −300 μV (% of all samples) | **$3.69 \times 10^{-2} \pm 7.74 \times 10^{-2}$** | $2.91 \times 10^{-2} \pm 6.32 \times 10^{-2}$ | $2.87 \times 10^{-2} \pm 5.92 \times 10^{-2}$ | $2.38 \times 10^{-2} \pm 5.00 \times 10^{-2}$ |
| −300 to −1000 μV (% of all samples) | **$2.87 \times 10^{-2} \pm 7.61 \times 10^{-2}$** | $2.45 \times 10^{-2} \pm 5.97 \times 10^{-2}$ | $2.09 \times 10^{-2} \pm 5.65 \times 10^{-2}$ | $1.30 \times 10^{-2} \pm 3.26 \times 10^{-2}$ |

| | 255-channel NS probe | | | |
|---|---|---|---|---|
| | Column 5 | Column 6 | Column 7 | Column 8 (Center) |
| −50 to −100 μV (% of all samples) | 0.422 ± 0.576 | 0.401 ± 0.561 | 0.383 ± 0.549 | 0.373 ± 0.547 |
| −100 to −200 μV (% of all samples) | 0.139 ± 0.263 | 0.134 ± 0.270 | 0.127 ± 0.269 | 0.119 ± 0.257 |
| −200 to −300 μV (% of all samples) | $2.37 \times 10^{-2} \pm 5.43 \times 10^{-2}$ | $2.08 \times 10^{-2} \pm 5.12 \times 10^{-2}$ | $1.80 \times 10^{-2} \pm 4.64 \times 10^{-2}$ | $1.43 \times 10^{-2} \pm 3.63 \times 10^{-2}$ |
| −300 to −1000 μV (% of all samples) | $9.36 \times 10^{-3} \pm 2.48 \times 10^{-2}$ | $5.13 \times 10^{-3} \pm 1.44 \times 10^{-2}$ | $3.64 \times 10^{-3} \pm 1.14 \times 10^{-2}$ | $4.45 \times 10^{-3} \pm 1.13 \times 10^{-2}$ |

**Supplementary Table 9.** Mean ± standard deviation of the amplitude distribution of the filtered potential in the four amplitude range for the first eight columns of recording sites of the 255-channel NeuroSeeker probe.

| | 255-channel NS probe | | |
|---|---|---|---|
| | Edge Left | Center | Edge Right |
| $-50$ to $-100$ μV (% of all samples) | $1.69 \times 10^{-2} \pm 2.62 \times 10^{-2}$ | $1.68 \times 10^{-2} \pm 2.14 \times 10^{-2}$ | **$2.37 \times 10^{-2} \pm 1.73 \times 10^{-2}$** |
| $-100$ to $-200$ μV (% of all samples) | $1.61 \times 10^{-3} \pm 2.29 \times 10^{-3}$ | $2.68 \times 10^{-3} \pm 4.48 \times 10^{-3}$ | **$5.56 \times 10^{-3} \pm 6.46 \times 10^{-3}$** |
| $-200$ to $-300$ μV (% of all samples) | $9.89 \times 10^{-5} \pm 1.56 \times 10^{-4}$ | $5.97 \times 10^{-4} \pm 1.22 \times 10^{-3}$ | **$6.62 \times 10^{-4} \pm 1.05 \times 10^{-3}$** |
| $-300$ to $-1000$ μV (% of all samples) | $2.74 \times 10^{-5} \pm 5.50 \times 10^{-5}$ | **$1.09 \times 10^{-4} \pm 2.76 \times 10^{-4}$** | $6.54 \times 10^{-5} \pm 1.03 \times 10^{-4}$ |
| No. of single units | **6.71 ± 5.02** | 5.00 ± 3.06 | **6.71 ± 2.93** |
| Amplitude of single units (μV) | 138.48 ± 66.29 | 153.98 ± 99.49 | **188.33 ± 118.54** |
| Isolation distance | 23.65 ± 18.52 | 33.35 ± 30.16 | **42.30 ± 48.72** |

**Supplementary Table 10.** Mean ± standard deviation of the calculated features for the 255-channel silicon probe data available online (www.kampff-lab.org/ultra-dense-survey).

| | 128-channel NS probe | |
|---|---|---|
| | Edge | Center |
| −50 to −100 µV (% of all samples) | **0.177 ± 0.193** | 0.162 ± 0.185 |
| −100 to −200 µV (% of all samples) | **$3.19 \times 10^{-2} \pm 4.65 \times 10^{-2}$** | $2.67 \times 10^{-2} \pm 4.07 \times 10^{-2}$ |
| −200 to −300 µV (% of all samples) | **$3.05 \times 10^{-3} \pm 5.52 \times 10^{-3}$** | $2.07 \times 10^{-3} \pm 4.44 \times 10^{-3}$ |
| −300 to −1000 µV (% of all samples) | **$8.45 \times 10^{-4} \pm 1.71 \times 10^{-3}$** | $5.33 \times 10^{-3} \pm 1.39 \times 10^{-4}$ |

**Supplementary Table 11.** Mean ± standard deviation of the amplitude distribution of the filtered potential in the four amplitude range for the larger dataset (n = 186 recordings) obtained with the 128-channel NeuroSeeker probe.

| | 128-channel NS probe | | | |
|---|---|---|---|---|
| | Edge Left | Center Left | Center Right | Edge Right |
| −50 to −100 μV (% of all samples) | **0.178 ± 0.197** | 0.164 ± 0.870 | 0.160 ± 0.184 | 0.177 ± 0.195 |
| −100 to −200 μV (% of all samples) | **$3.19 \times 10^{-2} \pm 4.91 \times 10^{-2}$** | $2.65 \times 10^{-2} \pm 4.06 \times 10^{-2}$ | $2.69 \times 10^{-2} \pm 4.18 \times 10^{-2}$ | $3.20 \times 10^{-2} \pm 4.78 \times 10^{-2}$ |
| −200 to −300 μV (% of all samples) | **$3.08 \times 10^{-3} \pm 6.20 \times 10^{-3}$** | $1.79 \times 10^{-3} \pm 4.32 \times 10^{-3}$ | $2.35 \times 10^{-3} \pm 5.17 \times 10^{-3}$ | $3.01 \times 10^{-3} \pm 5.89 \times 10^{-3}$ |
| −300 to −1000 μV (% of all samples) | **$9.21 \times 10^{-4} \pm 2.19 \times 10^{-3}$** | $4.74 \times 10^{-4} \pm 1.95 \times 10^{-3}$ | $5.93 \times 10^{-4} \pm 1.46 \times 10^{-3}$ | $7.67 \times 10^{-4} \pm 1.84 \times 10^{-3}$ |

**Supplementary Table 12.** Mean ± standard deviation of the amplitude distribution of the filtered potential in the four amplitude range for the large dataset (n = 186 recordings) obtained with the 128-channel NeuroSeeker probe, separated by left and right side.

| | 128-channel NS probe | |
|---|---|---|
| | Edge | Center |
| −50 to −100 μV (% of all samples) | **0.843 ± 0.511** | 0.816 ± 0.506 |
| −100 to −200 μV (% of all samples) | **0.255 ± 0.202** | 0.251 ± 0.212 |
| −200 to −300 μV (% of all samples) | **$3.45 \times 10^{-2} \pm 3.35 \times 10^{-2}$** | $2.94 \times 10^{-2} \pm 3.41 \times 10^{-2}$ |
| −300 to −1000 μV (% of all samples) | **$7.47 \times 10^{-3} \pm 7.63 \times 10^{-3}$** | $5.65 \times 10^{-3} \pm 7.88 \times 10^{-3}$ |

**Supplementary Table 13.** Mean ± standard deviation of the amplitude distribution of the filtered potential in the four amplitude range for the thalamic dataset (n = 9 recordings) obtained with the 128-channel NeuroSeeker probe.

| | 128-channel NS probe | | | |
|---|---|---|---|---|
| | Edge Left | Center Left | Center Right | Edge Right |
| −50 to −100 μV (% of all samples) | **0.853 ± 0.519** | 0.824 ± 0.510 | 0.808 ± 0.502 | 0.832 ± 0.510 |
| −100 to −200 μV (% of all samples) | **0.289 ± 0.242** | 0.265 ± 0.228 | 0.237 ± 0.197 | 0.218 ± 0.163 |
| −200 to −300 μV (% of all samples) | **$4.48 \times 10^{-2} \pm 4.93 \times 10^{-2}$** | $3.31 \times 10^{-2} \pm 4.03 \times 10^{-2}$ | $2.57 \times 10^{-2} \pm 2.80 \times 10^{-2}$ | $2.35 \times 10^{-2} \pm 1.98 \times 10^{-2}$ |
| −300 to −1000 μV (% of all samples) | **$1.04 \times 10^{-2} \pm 1.36 \times 10^{-2}$** | $6.92 \times 10^{-3} \pm 1.00 \times 10^{-2}$ | $4.37 \times 10^{-3} \pm 5.86 \times 10^{-3}$ | $4.62 \times 10^{-3} \pm 3.90 \times 10^{-3}$ |

**Supplementary Table 14.** Mean ± standard deviation of the amplitude distribution of the filtered potential in the four amplitude range for the thalamic dataset (n = 9 recordings) obtained with the 128-channel NeuroSeeker probe, separated by left and right side.

| | Neocorte× | | | |
|---|---|---|---|---|
| | Top | Upper Middle | Lower Middle | Bottom |
| −50 to −100 μV (% of all samples) | 0.201 ± 0.204 | 0.460 ± 0.271 | **0.638 ± 0.546** | 0.294 ± 0.344 |
| −100 to −200 μV (% of all samples) | $2.11 \times 10^{-2} \pm 2.85 \times 10^{-2}$ | $5.19 \times 10^{-2} \pm 4.54 \times 10^{-2}$ | **0.122 ± 0.143** | $6.15 \times 10^{-2} \pm 1.22 \times 10^{-1}$ |
| −200 to −300 μV (% of all samples) | $9.52 \times 10^{-4} \pm 1.39 \times 10^{-3}$ | $2.60 \times 10^{-3} \pm 3.20 \times 10^{-3}$ | $\mathbf{7.65 \times 10^{-3} \pm 1.01 \times 10^{-2}}$ | $5.22 \times 10^{-3} \pm 1.27 \times 10^{-2}$ |
| −300 to −1000 μV (% of all samples) | $2.39 \times 10^{-4} \pm 4.09 \times 10^{-4}$ | $1.19 \times 10^{-4} \pm 8.28 \times 10^{-5}$ | $\mathbf{4.79 \times 10^{-4} \pm 6.50 \times 10^{-4}}$ | $2.79 \times 10^{-4} \pm 5.91 \times 10^{-4}$ |

**Supplementary Table 15.** Mean ± standard deviation of the amplitude distribution of the filtered potential in the four amplitude range for the neocortical data obtained with the 128-channel probe, separated by longitudinal site position.

| | Thalamus | | | |
| --- | --- | --- | --- | --- |
| | Top | Upper Middle | Lower Middle | Bottom |
| −50 to −100 μV (% of all samples) | 0.714 ± 0.528 | 0.836 ± 0.541 | **0.929 ± 0.589** | 0.845 ± 0.634 |
| −100 to −200 μV (% of all samples) | 0.196 ± 0.213 | 0.248 ± 0.209 | **0.309 ± 0.262** | 0.283 ± 0.298 |
| −200 to −300 μV (% of all samples) | $2.15 \times 10^{-2} \pm 2.76 \times 10^{-2}$ | $2.39 \times 10^{-2} \pm 2.72 \times 10^{-2}$ | **$4.50 \times 10^{-2} \pm 5.26 \times 10^{-2}$** | $4.10 \times 10^{-2} \pm 5.74 \times 10^{-2}$ |
| −300 to −1000 μV (% of all samples) | $4.66 \times 10^{-3} \pm 5.70 \times 10^{-3}$ | $3.56 \times 10^{-3} \pm 4.80 \times 10^{-3}$ | **$9.88 \times 10^{-3} \pm 1.55 \times 10^{-2}$** | $9.52 \times 10^{-3} \pm 1.74 \times 10^{-2}$ |

**Supplementary Table 16.** Mean ± standard deviation of the amplitude distribution of the filtered potential in the four amplitude range for the thalamic data obtained with the 128-channel probe, separated by longitudinal site position.
